# CLASP2 stabilizes GDP-associated terminal tubulins to prevent microtubule catastrophe

**DOI:** 10.1101/2022.04.25.489454

**Authors:** Wangxi Luo, Vladimir Demidov, Qi Shen, Hugo Girão, Manas Chakraborty, Aleksandr Maiorov, Fazly I. Ataullakhanov, Chenxiang Lin, Helder Maiato, Ekaterina L. Grishchuk

## Abstract

CLASPs are ubiquitous stabilizers of microtubule dynamics but their molecular targets at the microtubule plus-end are not understood. Using DNA origami-based reconstructions we show that clusters of human CLASP2 form a load-bearing bond with terminal GDP-tubulins at the stabilized microtubule tip. This activity relies on the unconventional TOG2 domain of CLASP2, which releases its high-affinity bond with the GDP-dimers upon their conversion into polymerization-competent GTP-tubulin. By tethering dynamic microtubule ends near immobilized CLASP2, we show that the targets for CLASP2 binding at the polymerizing tip arise stochastically, leading to nanoscale disruptions in microtubule tip integrity. The ability of CLASP2 to recognize nucleotide-specific tubulin conformation and stabilize the catastrophe-promoting GDP-tubulins intertwines with the previously underappreciated exchange between GDP and GTP at terminal tubulins, providing a distinct molecular mechanism to suppress microtubule catastrophe without affecting tubulin incorporation.

## Introduction

Proper regulation of microtubule dynamics and load-bearing by microtubule tip-attached structures are essential for numerous cellular processes. CLASPs are an evolutionarily conserved family of ubiquitous factors that exert stabilizing effects on microtubules by suppressing catastrophes and promoting rescues (*1–4*). CLASPs importance has been recognized and extensively characterized in a variety of cellular and organismal systems, such as during cell migration, neuronal development, cytoskeleton reorganization in interphase and during cell division (reviewed in (*5, 6*)). CLASPs are highly potent regulators of the growing microtubule plus-ends, owing to their enrichment at these sites via the tip-binding proteins, such as EB. Microtubule ends are also regulated by localized multimolecular ensembles of CLASPs. For example, membrane associated clusters of CLASP molecules capture microtubule ends at the post synaptic membrane, whereas cortical clusters of CLASPs tether and stabilize microtubule ends near focal adhesions in motile cells (*7, 8*) or at cell cortex (*9–11*). These interactions are often modulated by cellular force, such as those produced during cell retraction (*12*), microtubule bundle protrusion into the lamella or contact-mediated cell–cell repulsion (*13*).

Studies in mitotic cells have been particularly informative for understanding the CLASP’s molecular mechanisms, because here CLASP proteins are clustered at kinetochores to mediate their dynamic interactions with microtubule plus-ends (*14–16*) Although kinetochore-microtubule attachments are subjected to considerable forces, the tethered microtubule ends are not static and undergo dynamic transitions. During metaphase, tubulin dimers are added continuously to the kinetochore-embedded microtubule ends, contributing to their poleward flux (Fig. 1A). Human cells encode two CLASP proteins, CLASP1 and CLASP2 (each with several isoforms), which play redundant roles in mitosis and assist poleward flux by promoting elongation of tethered microtubule ends under force (*15–18*)

**Figure 1.**
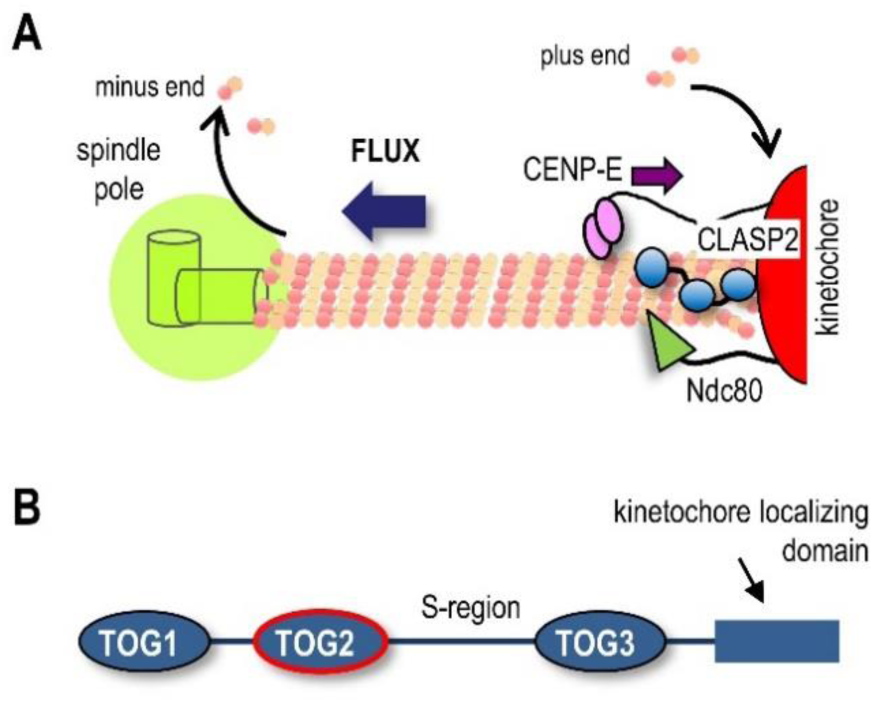
Kinetochore-associated structures and organization of human CLASP proteins. **A.** Schematic of kinetochore microtubule plus-end and its coupling to the kinetochore, which clusters several microtubule-binding proteins, including CLASPs, the Ndc80 proteins complex and the plus-end-directed kinesin, CENP-E. CLASP promotes incorporation of tubulins at the kinetochore-bound microtubule plus ends, sustaining poleward tubulin flux. **B.** Schematic of functional domains in human CLASP2α (not to scale).

The model in which CLASPs help to assemble tubulins at the kinetochore-embedded microtubule plus-ends has been attractive in view of the prior proposal that CLASPs act as a tubulin polymerase (*19, 20*). Specifically, a linked array of TOG domains (Fig. 1B), which are found in CLASPs and represent specialized tubulin-binding modules, have been thought to be instrumental in promoting tubulin dimers incorporation (*21–23*). However, recent experiments in vitro have found that a single TOG2 domain, supplemented with the downstream flanking region of human CLASP2 (TOG2-S protein) is sufficient to provide anti-catastrophe and rescue activities characteristic of the full length proteins (*24, 25*).

TOG2 is unusual among the set of all TOG domains, as its highly bent shape is not compatible with binding to straight tubulins in the microtubule wall. Curvature of TOG2 also exceeds the curvature of the crystallized forms of soluble tubulin (*26, 27*). Accordingly, CLASPs have been proposed to bind to highly curved protofilaments at the microtubule tip (*28*). High-resolution fluorescence microscopy, however, localizes TOG2-S to a region trailing the growing microtubule tip by about 90 nm (10-12 tubulin layers), leading to a proposal that this is the actual site of its anti-catastrophe activity (*24*). Thus, the primary molecular target for TOG2 binding at the growing microtubule end, and the mechanism of its stabilizing activity, are ill-understood. One underexplored model for CLASPs microtubule binding posits that it is regulated directly by GTP, based in part on the presence of molecular features of a GTP-binding site in the CLASP orthologue in Drosophila (*3*). Whether this putative GTP-binding site plays an important role in CLASP2 activity in vitro or in mitotic cells remains unknown.

Because of the pivotal role of CLASPs in numerous cellular functions, we have sought to gain insights into its role in regulating dynamics of the tethered microtubule ends. With the DNA origami technique, we show that microtubule plus-ends bound to clusters of CLASP2 are not capable of elongation, arguing against the direct role of CLASP2 in promoting tubulin incorporation. We reconstructed high-affinity end-association using only the TOG2 domain with flanking regions and found that microtubule end is released by soluble GTP independently of the putative GTP-binding site. Stoichiometry of CLASP2 binding to stabilized microtubule tip, as well as its marked GTP-sensitivity, strongly suggest that TOG2 targets naturally occurring terminal GDP-tubulins, rather than tubulin protofilaments or tubulins at a tip-proximal location. We therefore proposed a model in which CLASP2 promotes microtubule elongation indirectly, by stabilizing the catastrophe-promoting terminal GDP-tubulin dimers, which are generated stochastically during GTP-tubulin assembly.

## Results

### DNA origami provides a multivalent molecular platform to reconstruct physiological clusters of kinetochore-associated proteins

To seek insight into the molecular mechanisms of CLASP2 activity we used DNA origami technology (*29, 30*), which provides a versatile platform for organizing protein molecules with precisely controlled stoichiometry and spatial positioning (*31, 32*). Specifically, we modified a previously developed DNA-origami nano-circle (*33*) to function as the structural scaffold for organizing kinetochore-associated proteins. Its 62 nm outer diameter is a good match for the overall dimensions of one microtubule-binding site at the kinetochore (Fig. 2A), which tethers flared tubulin protofilaments of the microtubule plus-end (*34*). This scaffold features three types of ssDNA handles (Supplementary Fig. 1, see Methods). Eight handles originating from the inner surface of the circular core were hybridized with complementary ssDNAs labeled with Cy5 to assist scaffold visualization. Four biotin-labeled anti-handles were used to firmly flatten these circles on neutravidin-coated glass slides. Next, ssDNAs functionalized with benzyl-guanine (BG) were covalently attached to a protein adaptor consisting of a SNAP-tag (New England Biolabs, Inc.) fused to GBP, a GFP-binding nanobody (*35*) SNAP-GBP-containing anti-handles were annealed to the 24 ssDNA handles on the outer surface of the DNA origami scaffold (Fig. 2B), enabling high-affinity recruitment of any soluble GFP-tagged protein that was brought into a microscopy chamber. After unbound protein was washed away, over 95% of the Cy5-labeled scaffolds colocalized with bright GFP dots, implying formation of high-density molecular clusters (Fig. 2C).

**Figure 2.**
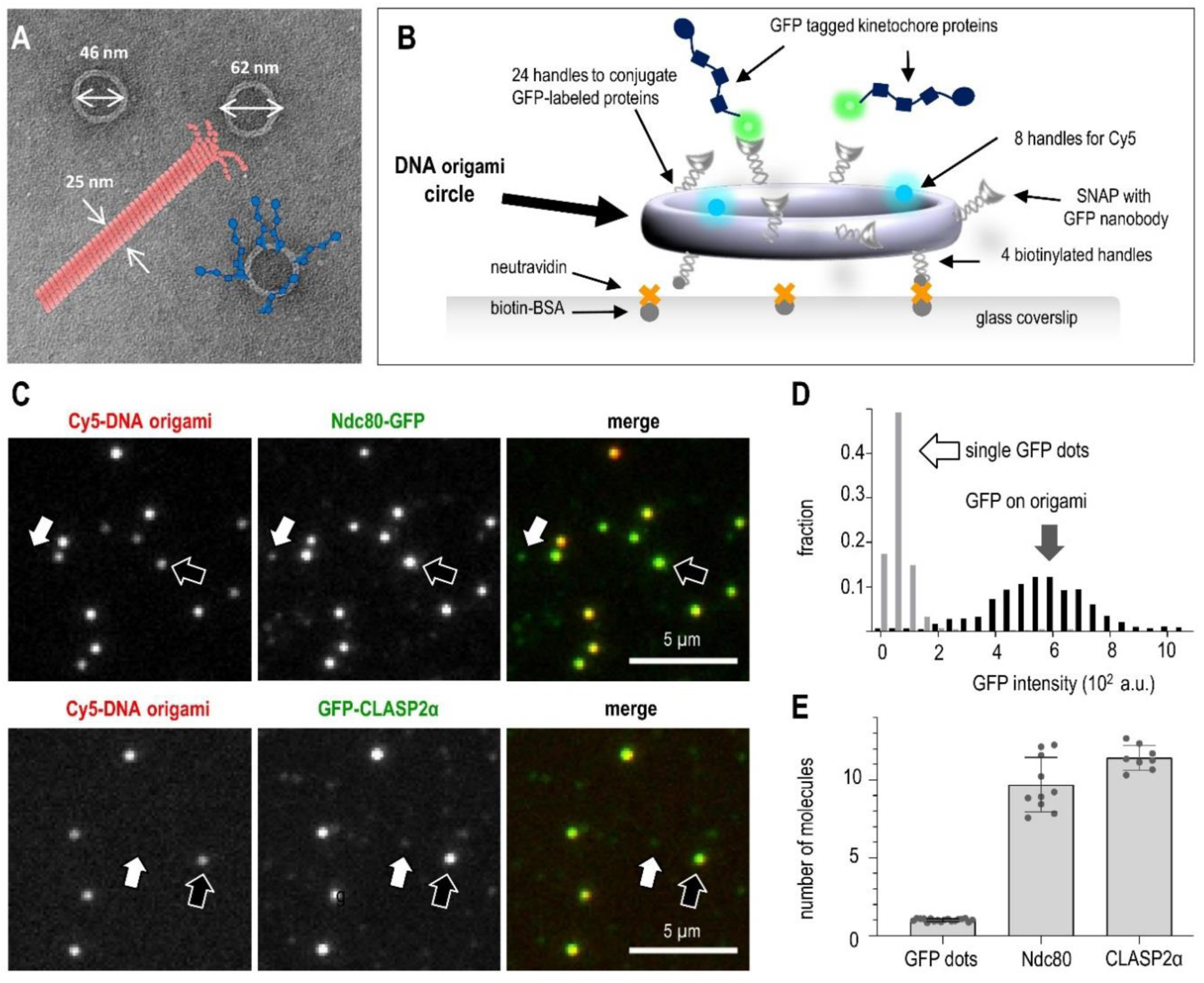
Reconstruction of multivalent molecular clusters for kinetochore-microtubule interaction studies. A. Electron microscopy image of the DNA origami circles with overlaid schematics of a microtubule (drawn to scale), and recruited kinetochore proteins (blue). B. The use of an immobilized DNA scaffold and conjugate GFP-tagged proteins to make a microtubule-binding site. C. Representative fluorescence images of Cy5-labeled DNA scaffolds (examples indicated by black arrows) with conjugated GFP-tagged proteins: human Ndc80 Broccoli and full length CLASP2α. White arrows point to dim dots with GFP but not Cy5 fluorescence, representing single GFP-tagged protein molecules that attach non-specifically. D. Typical GFP intensity distribution of DNA scaffolds (n = 180) and dim dots (n = 527) based on one representative experiment with Ndc80-GFP, in which 13 image fields were analyzed. E. Number of GFP-tagged molecules per DNA origami scaffold. Each dot represents mean of the GFP intensity distributions from independent experiments plotted relative to the mean intensity of single GFP molecules. Error bars are SEM.

Dim GFP dots were also seen non-specifically attached to the coverslip. Their uniform brightness and single-step photobleaching kinetics indicated that they represented single molecules (Supplementary Fig. 2, see Methods). By dividing average GFP intensity of the scaffold-associated protein clusters by the single molecule brightness, we estimate that the clusters recruit on average 10-12 protein molecules (Fig. 2 D,E), which is a reasonable approximation of a microtubule-binding site at a kinetochore (*36*). Thus, the generated molecular clusters include a sufficient number of each protein under study to engage in multivalent interactions with one microtubule end. As such, they represent physiologically-relevant assemblies whose study in vitro may elucidate kinetochore-microtubule interactions in vivo.

### Molecular clusters of CLASP2 but not Ndc80 protein form durable and specific attachment to stabilized microtubule plus-ends

Previous in vitro assays using soluble CLASP proteins revealed their diffusion on microtubule walls, as well as slight enrichment at the polymerizing microtubule plus-ends (*20, 24, 37, 38*). To investigate the microtubule-binding properties of the origami-clustered assemblies of CLASP molecules, we used full length human CLASP2α, which is among the most studied CLASP isoforms. Taxol-stabilized microtubules labeled with rhodamine were flowed into the chambers with coverslip-immobilized clusters of CLASP2α and imaged using Total Internal Reflection Fluorescence (TIRF) microscopy (Fig. 3A, see Methods). Most of the microtubules associated with CLASP2α-containing clusters appeared as blurry spots, suggesting that tethered microtubules orient away from the coverslip-surface (Video 1). In contrast, clusters containing a different kinetochore protein, the microtubule-wall diffusing Ndc80 complex, recruited microtubules by their walls, and near the microtubule tip (Fig. 3B). These microtubules were often seen diffusing on the coverslip-bound Ndc80-clusters while remaining within the TIRF imaging plane (Video 2) (Supplementary Fig. 3).

**Figure 3.**
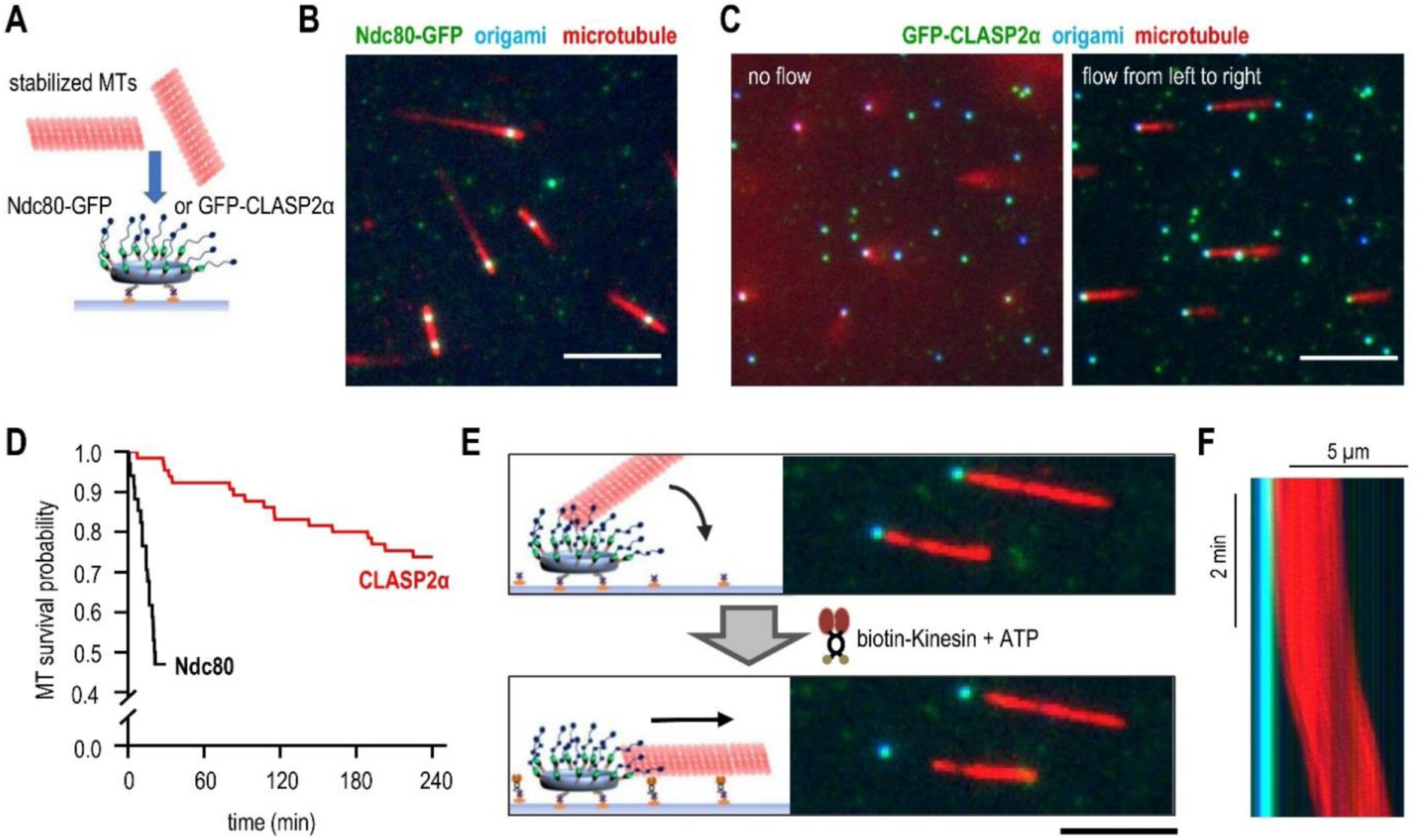
End-on attachment of microtubules to molecular clusters of full length CLASP2α protein. A. Schematic of experimental assay with a DNA origami circle attached to a coverslip and presenting clusters of microtubule-binding proteins to rhodamine-labeled taxol-stabilized microtubules (MTs). B. Representative TIRF image of microtubules (red) bound to DNA origami scaffolds (blue) with clusters of GFP-tagged Ndc80 Broccoli (green). Bar is 5 µm. C. Same as in panel (B) but with CLASP2α clusters and showing the same field with (right) or without (left) buffer flow. D. Lifetime of microtubule association with the clusters of the indicated proteins. Kaplan–Meier survival plots were built based on n microtubules observed in N independent experiments: for Ndc80 n=34, N=2; for CLASP2α n=65, N=3. E. Schematic of the experiment and representative images of microtubules initially tethered at CLASP2α clusters, but then dragged away by the coverslip-bound, plus-end-directed kinesin K560 in the presence of 10 µM ATP. Bar is 5 µm. F. Kymograph illustrating the lag period after addition of ATP and the persistence of the microtubule gliding away from the CLASP2α cluster (overlaid blue, green and red images, see panel C for details). Variations in red intensity are due to uneven illumination.

Such diffusive motions on the microtubule surface were notably absent when CLASP2α clusters were used. To reveal the preferred mode of microtubule-binding by the CLASP2α clusters, we applied a gentle flow of imaging buffer (Video 3). Under the flow force, the scaffold-associated microtubules came into focus, revealing one end of each microtubule tethered to a CLASP2α cluster (Fig. 3C). These attachments were very durable, as half of microtubules maintained end-association for 4 hours, significantly longer than the microtubule-binding of Ndc80 clusters of similar size (Fig. 3D).

To determine whether the tethered microtubules had a specific polarity, we took advantage of the plus-end-directed Kinesin 1. After the microtubules formed end-on attachment to the CLASP2α clusters, we conjugated kinesin motors to the coverslip surface and introduced a flow of imaging buffer supplemented with Mg-ATP (Fig. 3E). Microtubules pivoted at the CLASP2α-tethered ends, landed on the coverslip, and after a brief delay began to glide unidirectionally (Fig. 3F, Video 4). All microtubules examined (n=124) glided away from the tethering sites, not toward them. Because kinesin propels microtubules with the minus-end leading, these findings reveal that clusters of CLASP2α form durable and highly specific attachment to the microtubule plus-ends, same polarity of microtubule binding seen at kinetochores.

### Microtubule ends tethered to the CLASP2 clusters can withstand significant load

Microtubule ends embedded at kinetochores of dividing cells can withstand significant forces (*39–42*). To investigate the ability of CLASP2α clusters to maintain association with the stabilized microtubule plus-ends under load, we used a laser trapping approach (see Methods, Supplementary Fig. 4). Polystyrene beads coated with anti-tubulin antibodies were flowed into the chamber, and they attached spontaneously to the walls of microtubules whose ends were tethered to the coverslip-immobilized clusters of CLASP2α (Fig. 4A,B). A microtubule-bound bead was then captured in a stationary optical trap and the stage was moved at a constant speed to pull the cluster away from the laser-trapped bead. Because CLASP2α-tethered microtubules project into the chamber, bead displacement was monitored in two directions: along the stage motion (*y*-axis) and along the *z*-axis, pointing away from the coverslip.

**Figure 4.**
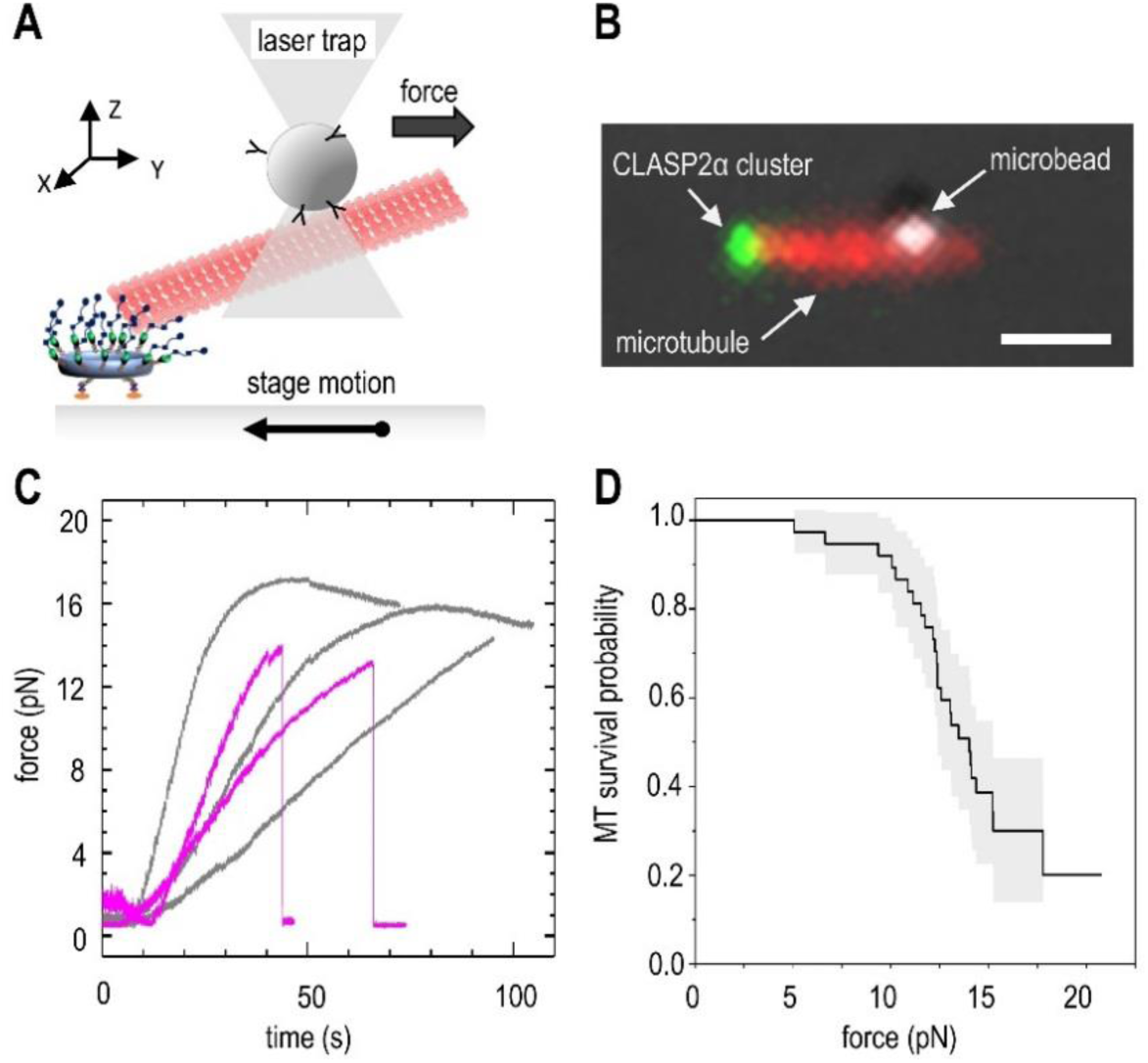
Measuring the strength of a microtubule end-on attachment using laser trapping. A. Schematic of the experimental assay in which a pulling force is applied to the microtubule tethered at its plus-end to a coverslip-immobilized CLASP2α cluster. Microtubule is held in the laser trap via the wall-bound microbead while the stage is moved in the opposite direction until a bond is ruptured or the maximum force is reached with no rupture. B. Representative fluorescence image of a taxol-stabilized rhodamine-labeled microtubule and the coverslip-bound CLASP2α cluster overlaid with the DIC image of the microtubule-bound bead. Because the bead is located several microns away from the coverslip (deep into the chamber), the microtubule image represents a projection, constructed by combining several images taken at different z planes. Bar is 1 μm. C. Force application curves from individual experiments. Two typical outcomes are illustrated by these five curves. Grey curves show increase in the pulling force followed by a gradual decrease, indicating that a maximal force was reached with no rupture. Curves in purple show similar increases in pulling force, followed by an abrupt drop, indicative of a rupture. D. Kaplan-Meier survival plot based on 27 experiments. Grey area corresponds to 95% confidence interval.

Using this approach, we detected a rupture in this mechanical system in 26 out of 39 experiments. Rupture was evident via an abrupt decrease in force exerted by the laser on the trapped bead (Fig. 4C, purple). In 25 of these events, the beads remained bound to the microtubule and GFP signal from CLASP2α persisted at the same location on the coverslip (Supplementary Fig. 4D), indicating that the rupture occurred specifically between CLASP2α and the microtubule plus-end. In only one case, the rupture took place between the origami and the coverslip. The remaining microtubules (n=13) did not detach from the CLASP2α clusters even under highest pulling force our laser trap could apply: 15.2 ± 2.8 pN (Fig. 4C, grey). Kaplan-Meier survival plot in Figure 4D shows that the 50% survival probability for CLASP2α attachment to microtubule plus-end is 14 pN (95% confidence interval: 12.4 - 15.2 pN). These results strongly suggest that a cluster of 10-12 CLASP2α molecules is able to provide a significant contribution to microtubule attachment stability and to mediate force-dependent cellular functions, for example, at kinetochore-tethered microtubule ends during chromosome segregation.

### Soluble GTP destabilizes the end-on microtubule attachment to CLASP2

Next, we investigated whether clusters of CLASP2α can promote addition of soluble tubulins to tethered microtubule plus-ends; such tubulin incorporation occurs during kinetochore microtubule flux (Fig. 5A). To facilitate tubulin polymerization, we used microtubule seeds polymerized with GMPCPP. The seeds formed end-on attachment to the CLASP2α clusters similarly to the taxol-stabilized polymers. When soluble GTP-tubulin was added into the chamber, no tubulin addition was seen at the tethered ends. Instead, the end-bound microtubules detached immediately (Fig. 5B). We ruled out that our tubulin or buffer composition did not support microtubule elongation by observing in the same chambers that tubulin assembly was possible at the ends of rare microtubules that bound laterally to the CLASP2α clusters (Video 5). Thus, these results reveal unexpectedly that CLASP2α binding to the microtubule end is incompatible with tubulin addition to that end.

**Figure 5.**
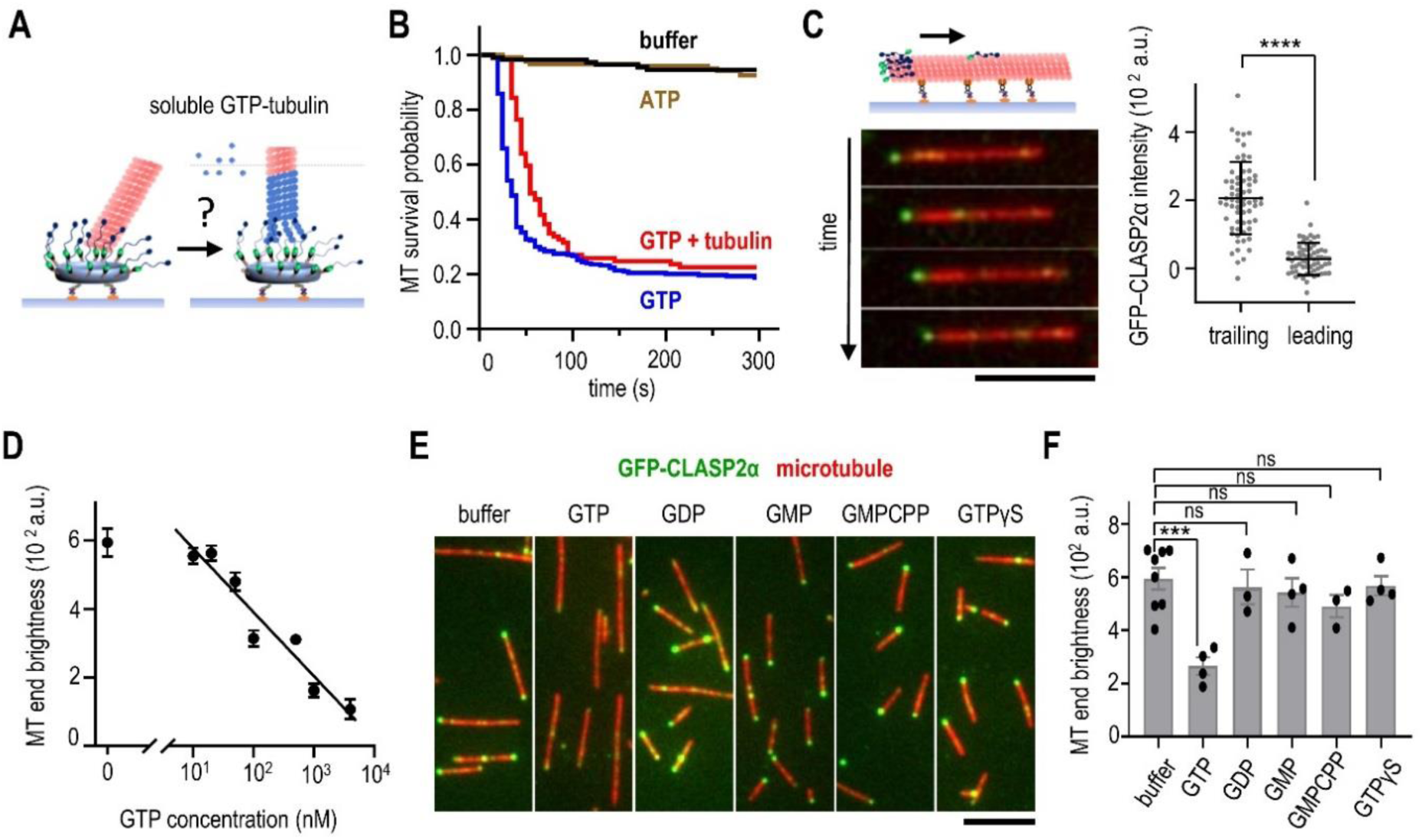
Soluble GTP as a highly specific inhibitor of microtubule end-on attachment to CLASP2α. A. Schematic of the experiment to examine ability of CLASP2α clusters to incorporate soluble tubulin (blue) at an associated microtubule plus-end (red). B. Lifetime of microtubule end-on attachments to CLASP2α clusters. Kaplan-Meier plots show the probability of attachment survival after addition of imaging buffer, buffer with 1 mM Mg-ATP, with 1 mM Mg-GTP or with soluble tubulin (4 μM tubulin with 1 mM Mg-GTP). Curves are based on n microtubules imaged in N independent experiments: buffer n=110, N=2; ATP n=120, N=2; GTP n=255, N=4, tubulin+GTP n=89, N=3. C. Time-lapse images of a taxol-stabilized rhodamine-labeled microtubule gliding on coverslip coated with Kinesin 1 in the presence of 1 nM soluble GFP-tagged CLASP2α and 1 mM Mg-ATP. Bar is 5 µm. Graph on the right shows CLASP2α brightness at the trailing (minus) and leading (plus) microtubule ends: mean ± S.D., n=66 microtubules, N=2. Significance was determined using Paired t test. **** p < 0.0001. D. Inhibition curve for soluble GFP-tagged CLASP2α binding to taxol-stabilized microtubule ends plotted on a semi-log scale and with linear fitting. Dots correspond to the average brightness of the microtubule end-associated GFP-CLASP2α signal determined in the presence of indicated concentration of Mg-GTP (> 30 microtubules for each point). Error bars are standard deviation. E. Representative images of taxol-stabilized microtubules incubated with 1 nM soluble GFP-tagged CLASP2α in the presence of 1 µM of indicated nucleotides. F. Intensity of CLASP2α signal at the brightest microtubule end in the presence of 1 µM of indicated nucleotides. Each dot represents the average intensity of at least 50 microtubule ends collected in an independent experiment (see Data Source file for details). Error bars are SEM, Significance was determined using Welch’s t test. *** p < 0.001; ns – not significant.

To tease apart this relationship we varied experimental conditions and discovered that GTP alone, rather than GTP-tubulin, leads to a precipitous loss of end-on attachment by CLASP2α (Fig. 5B). The addition of ATP does not destabilize microtubule end-attachment, suggesting that the effect is nucleotide specific. Because the close proximity of microtubule-binding proteins to the charged DNA nano-circle might affect interactions between the origami-conjugated proteins and the microtubule end (*43*), we next asked whether GTP-dependent CLASP2α dissociation was dependent on DNA scaffolds. Taxol-stabilized microtubules were immobilized on the coverslip using tubulin antibodies and visualized via TIRF microscopy. When soluble GFP-tagged CLASP2α was added to this preparation in a GTP-free buffer, one end of most microtubules became obviously fluorescent, with weaker spots seen along microtubule walls and at the opposite microtubule end. Using microtubule gliding assay with Kinesin 1 and soluble ATP (Fig. 5C), we observed that 95% of bright microtubule ends with GFP-CLASP2α were trailing (N = 66 microtubules; Video 6). Thus, a preferred localization of CLASP2α to the plus-ends of stabilized microtubules does not depend on DNA origami scaffolds.

Using the origami-free system, we investigated the impact of soluble GTP on microtubule end decoration by CLASP2α and found that it inhibits the reaction in a concentration dependent manner with IC_50_ = 320 nM (Fig. 5D, Supplementary Fig. 5). This effect was highly specific to GTP, as other guanosine-containing nucleotides, such as GDP and GMP, and the non-hydrolyzable GTP analogs GMPCPP and GTPγS, had virtually no effect on interactions between CLASP2α and microtubule plus-ends (Fig. 5 E,F). Together, the results using soluble and clustered CLASP2α strongly suggest that GTP causes a rapid and highly specific dissociation of CLASP2α from the microtubule plus-end.

### Putative GTP-binding site in the TOG2 domain of CLASP2 is not essential for microtubule interactions in vitro and in mitotic cells

The GTP-sensitivity of binding between a stabilized microtubule end and CLASP2 is surprising. We were intrigued by this finding in view of a prior work reporting that microtubule binding by the *Drosophila* CLASP orthologue is regulated by GTP (*3*). Indeed, this prior work identified sequences in *Drosophila* CLASP that match putative GTP-binding motifs: the NKLD sequence and Walker A-like sequence GGGTGTG (*44, 45*). Sequence alignment of CLASP2 proteins from different species revealed a high degree of conservation of NKxD sequence, with x represented by phenylalanine in human CLASP2 (Supplementary Fig. 6). However, there was no obvious Walker A sequence at the same location as in *Drosophila* CLASP or in other parts of CLASP2α. Because the amino acid composition of the GTP-binding pockets in different proteins is diverse, these alignments do not rule out that human CLASP2α can be regulated directly by GTP. Interestingly, the conserved NKxD motif localizes to the TOG2 domain, which is responsible for the microtubule-stabilizing activity of CLASP2 (*6*). We therefore hypothesized that 1) the TOG2 domain is involved in GTP-dependent microtubule-end binding, and 2) this regulation requires the NKFD sequence.

First, we constructed L-TOG2-S protein in which the globular TOG2 domain is flanked by disordered L and S regions (Fig. 6A). Bacterially expressed and purified GFP-tagged L-TOG2-S readily decorated one end of taxol-stabilized microtubules (end brightness 594 ± 41 a.u. for full length protein vs. 757 ± 57 a.u. for L-TOG2-S protein), consistent with this domain’s importance for CLASP2α binding to microtubule plus-ends. This association was reduced in the presence of 1 μM GTP but not GMPCPP (Fig. 6 B,C), revealing a nucleotide sensitivity similar to that of CLASP2α and suggesting that this domain is responsible for the GTP-sensitivity of the whole protein’s microtubule binding.

**Figure 6.**
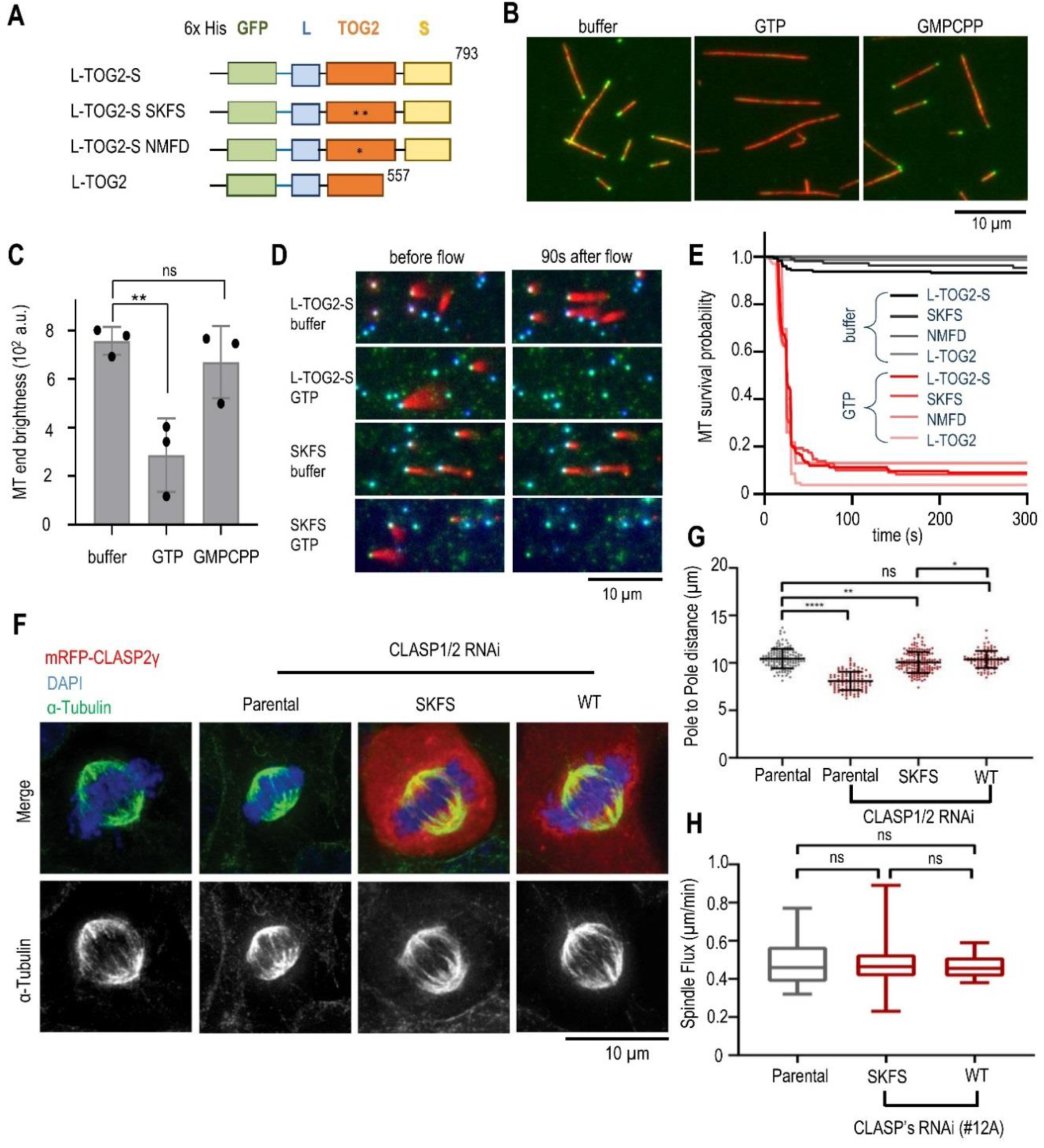
Probing the role of putative GTP-binding site in TOG2 domain. A. Schematics of truncated CLASP2α constructs used in this work. SKFS mutant protein has two substitutions N403S and D406S, NMFD mutant has one substitution K404M. All proteins were tagged with N-terminal GFP, then expressed and purified as described in Methods. Numbers indicate amino acids in wild type protein sequence. Asterisks indicate introduced mutations. B. Representative TIRF images of taxol-stabilized microtubules (red) incubated with 1 nM GFP-L-TOG2-S. Decoration of microtubule ends is visible in the buffer with no nucleotide or with 1 µM GMPCPP, but not in the presence of 1 µM GTP. C. Intensity of GFP-L-TOG2-S signal at microtubule end. Each dot represents average intensity from at least 50 microtubule ends observed in an independent experiment (see Data Source file for details). Error bars are SEM. D. Still images from video recording of experiments in which indicated proteins were conjugated to the coverslip-immobilized DNA scaffolds (left images). Images on the right were taken after the addition of buffer with or without 1 mM GTP. E. Kaplan–Meier survival plots for microtubule association with different protein clusters in the buffers with or without 1 mM GTP, see Data source file for details. F. Immunofluorescence analysis of spindle length in metaphase in different U2OS PA-GFP-α-tubulin cell lines expressing the mRPF-CLASP2γ constructs with (SKFS) or without (wild type) mutation of the NKFD motif. Scale bar is 5 µm. G. Quantification of mitotic spindle length: the first column represents the parental U2OS cell line and the remaining columns represent treatment with RNAi for both endogenous CLASP1 and CLASP2. Each data point represents an individual cell. Bars represent mean and standard deviation. Quantifications from 3 independent experiments: Parental Scramble RNAi= 133 cells; Parental CLASPs RNAi = 105 cells; SKFS mutant CLASPs RNAi = 155 cells; Wild Type CLASPs RNAi = 90 cells. ns = non-significant, * = p<0.05, ** = p<0.01, **** = p<0.0001, t-test. H. Quantification of spindle poleward flux. The first column corresponds to the parental cell line, and the remaining columns correspond to mRPF-CLASP2γ expressing cell lines after endogenous CLASP1 and CLASP2 depletion. Boxes represent median and interquartile interval; bars represent minimum and maximum values. Parental = 10 cells, pool of 3 independent experiments; SKFS mutant = 24 cells, pool of 5 independent experiments; Wild Type = 8 cells, pool of 3 independent experiments. ns = non-significant, Mann-Whitney U-Test.

To test whether GTP acts by binding to the putative GTP-binding site in TOG2, we replaced this site with the amino acid sequences NMFD and SKFS. When conjugated to DNA scaffolds, wild type L-TOG2-S and its mutant versions, as wells as a shorter truncation of wild type protein L-TOG2, were able to tether microtubule ends. Microtubule ends released rapidly upon addition of soluble GTP by all these constructs (Fig. 6 D,E). Thus, although the TOG2 domain appears to be the primary molecular determinant of GTP-sensitive microtubule end-on affinity, this interaction in vitro is not dependent on the NKFD sequence.

We then worked to discover any functional relevance of NKFD for mitosis in human cells. We engineered a mutant CLASP2 sequence with SKFS and tested whether it can rescue RNAi knockdown of endogenous CLASP1 and 2 in U2OS cells (see Methods. In a recent study (*18*) it was shown that the smallest isoform of hCLASP2, CLASP2γ is sufficient to ensure all the normal functionalities of both CLASP1 and CLASP2 at the kinetochore-microtubule interface, indicating that the TOG1 domain of CLASP2α is dispensable for mitosis. In addition, it was also previously reported that disruption of CLASPs function in mitotic cells severely compromises soluble tubulin incorporation at the plus-ends of kinetochore-bound microtubules, causing shortening of metaphase spindles (*14, 15, 46*), so we examined this phenotype in cells expressing the SKFS mutant. We found that the mean metaphase spindle length decreased very little in these cells: from 10.5 μm in the wild-type rescue to 10 μm (Fig. 6 F,G). If GTPase activity were essential for a CLASP2 role at the kinetochore-microtubule interface, we would have expected a much stronger spindle shortening phenotype in the SKFS mutant, comparable with the disruption of the TOG2 domain (*18*). As a direct assessment of how the SKFS impacts the rate of tubulin incorporation at the kinetochore-embedded microtubule plus-ends, we measured the rate of poleward flux in these cells, but it was also unchanged (Fig. 6H). Using fluorescence dissipation after photoactivation, we then examined microtubule turnover rate. This approach did not reveal any significant difference in the half-life of the kinetochore and non-kinetochore microtubules between parental cells and cells rescued with wild-type or SKFS mutant proteins (Supplementary Fig. 7). We concluded that the NFKD motif is dispensable for CLASP2’s role at the kinetochore-microtubule interface.

### CLASP2 binds specifically to terminal tubulins, which have exchangeable GDP-GTP binding pockets

To seek other molecular explanation for the GTP-sensitivity of CLASP2α affinity to microtubule plus-ends, we considered an alternative hypothesis in which the GTP dependency is mediated by polymerized tubulins, rather than CLASP2α. Both, α- and β-tubulins have GTP binding sites, but within the microtubule wall, GTP nucleotide is associated only with α-tubulin (N site), whereas the β-tubulin’s “E-site” is occupied by GDP (*47*). Thus, soluble GTP might exert its effect on stabilized microtubules in our assays by interacting with the E-sites in β-tubulins (Fig. 7A). Although there is no evidence of a significant GTP binding to this site within the microtubule wall, the GTP-binding site at the most distal β-tubulin is exposed and permits nucleotide exchange (*48*).

**Figure 7.**
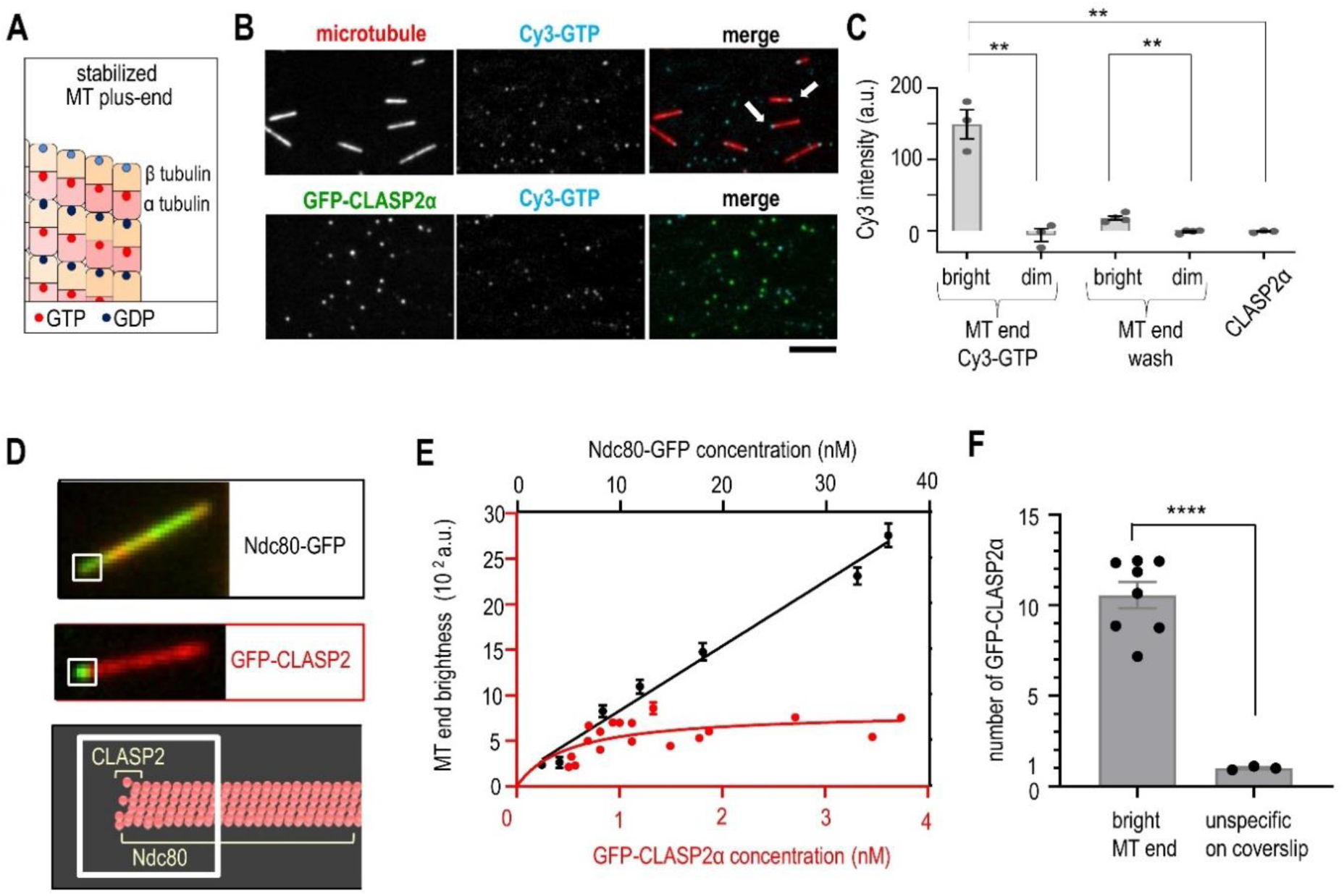
GTP and CLASP2α binding to terminal tubulins at stabilized microtubule tips. A. Schematic of the plus-end of stabilized microtubule showing GDP (dark blue) associated with β-tubulins and GTP (red) associated with α-tubulin. The E-sites of terminal tubulins are shown in light blue. B. Representative TIRF images of taxol-stabilized microtubules (upper row) and DNA origami-based clusters of full length CLASP2α (lower row) incubated with 100 nM Cy3-GTP. Some Cy3-GTP molecules bind non-specifically to the coverslips, but only microtubule images show colocalization (arrows point to example microtubule tips with prominent Cy3 signal). Images in the Cy3 channel were taken and processed using same settings and procedures, allowing direct comparison of signal brightness at microtubule tips and on CLASP2α clusters. Bar 5 µm. C. Cy3 intensity of microtubule ends and of CLASP2α clusters incubated with 100 nM Cy3-GTP, as in panel B. “MT end wash” images were captured after Cy3-GTP was replaced with imaging buffer. Each dot represents average brightness from an independent experiment, see Data Source file for details. Error bars are standard error of the mean, ** = p<0.01, t-test. D. Example images of taxol-stabilized microtubules decorated with GFP-labeled CLASP2α or Ndc80 Broccoli. Brightness was determined at the microtubule ends using same size regions (white squares) for both proteins, so the maximum brightness reflects the number of binding sites for these proteins (schematic). E. GFP intensity of microtubule ends decorated with proteins as in panel D. For easier comparison, data for different proteins are plotted using different scales. Each dot represents the median intensity of at least 32 microtubule ends, error bars are SEMs, see Data Source file for statistics. Lines are linear regression (black) or hyperbolic fit (red) with *K_d_* = 0.5 ± 0.4 nM and plateau *I_max_* = 800 ± 200 a.u. F. Normalized Intensity of microtubule ends decorated with 1 nM GFP-CLASP2α and the GFP-CLASP2α dots bound non-specifically to the coverslips in the same experimental chambers and imaged using same settings and protocols. Each point on the graph represents average intensities for one experiment trial with total number of microtubule ends n=718 and non-specific dots n=305. To normalize, average single molecule intensity was assumed to equal 1.

We tested whether soluble GTP can bind to the microtubule tips under our assay conditions by using Cy3-labeled GTP (*49*). TIRF imaging revealed that Cy3-GTP highlighted the ends of taxol-stabilized microtubules, with one end showing much brighter intensity (Fig. 7B). In contrast, clusters of full length CLASP2α protein incubated with Cy3-GTP and imaged under identical conditions showed no Cy3 signal (Fig. 7 B,C). Thus, microtubule ends but not CLASP2α are the primary receptor for GTP in our assays, consistent with our conclusion from the mutational analysis of CLASP2α. The Cy3-GTP signal at the microtubule ends was reduced by washing with a nucleotide-free buffer (Fig. 7C), implying that this binding is reversible. Moreover, fluorescently labeled GTP was able to reduce the intensity of CLASP2α end-decoration (by ∼1.9-fold at 100 nM, N = 186 microtubule ends), confirming that this fluorescently labeled nucleotide is functionally active. Thus, the E-site of the terminal β-tubulin in taxol-stabilized microtubules appears to have no nucleotide in our assay buffers, but it readily accepts and releases soluble GTP.

We hypothesized that GTP binding to the terminal tubulin dimer is a direct cause of CLASP2α dissociation. If this model is correct, there should be no more than 13 binding sites for CLASP2α at the microtubule tip, which we tested using soluble GFP-tagged CLASP2α and taxol-stabilized microtubules (Fig. 7D). Microtubule end brightness increased gradually with increasing concentration of CLASP2α and plateaued, consistent with saturation of its binding sites (Fig. 7E). In contrast, soluble GFP-tagged Ndc80 decorated the same area at the microtubule tips to a higher degree due to a vastly greater number of binding sites (Supplementary Fig. 8), demonstrating that saturation of the CLASP2α signal is not caused by a technical limitation of our quantitative procedure. By comparing the maximum brightness of the CLASP2α tip-decoration with the intensity of single molecules imaged in the same chamber, we estimate that end-binding plateaued at 12±2 molecules of CLASP2α (Fig. 7F). This number matches closely the number of microtubule protofilaments, and therefore the number of terminal β-tubulins with exposed GTP-binding pockets.

### CLASP2 softens occasionally the growing microtubule tip

Next, we sought to gain more insight into the activity of CLASP2α clusters at dynamic microtubule tips. Because tight CLASP2α binding to the terminal dimers at the microtubule tip is incompatible with tubulin incorporation, another protein must assist tethering of the microtubule end near the immobilized CLASP2α cluster. We took advantage of our previously developed strategy to reconstruct tethering of dynamic microtubule ends with help from a plus-end-directed kinesin (*50*). Brightly-labeled GMPCPP microtubule seeds were allowed to bind to coverslip-immobilized beads coated with a mixture of full length CLASP2α and the motor domains of CENP-E kinesin, which in mitotic cells recruits CLASPs in a motor-independent manner and promotes close proximity between microtubule plus-ends and kinetochores(*15*) (Fig. 1A). ATP was added and CENP-E motor drove the microtubules to bring their plus-ends closer to bead’s surface (Fig. 8A). Unlabeled GTP-tubulin was then added and tubulin incorporation at the plus ends was monitored via the motion of the labeled seeds away from the beads.

**Figure 8.**
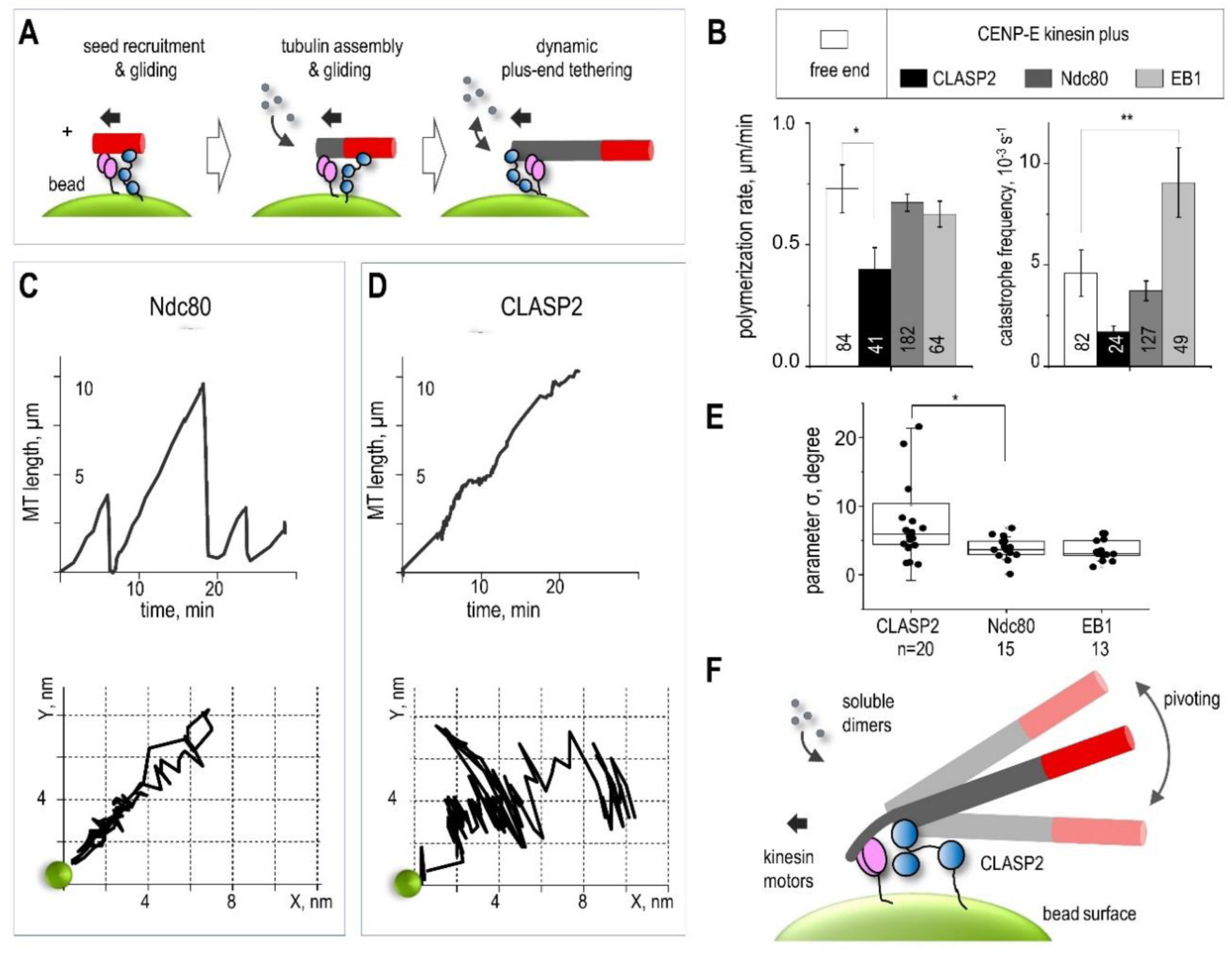
Dynamics of the microtubule end tethered near bead-immobilized CLASP2α and comparison with other proteins. A. Schematic of the experimental assay in which initial lateral attachment of the GMPCPP-stabilized microtubule seed is converted into the end-on attachment via the combined activity of the bead-conjugated CENP-E kinesin and a microtubule wall-diffusing protein. Black arrow shows the direction of the walking motor, which drives microtubule gliding in opposite direction until the motor reaches the microtubule plus-end. In the presence of soluble GTP-tubulin, gliding becomes coupled with microtubule dynamics at the bead-tethered end. Dynamics of this end and its mechanical rigidity are deduced from the directional and sideways motions of the labeled seed (in red). B. Dynamic parameters of the plus end of freely growing microtubules and of the plus-ends coupled to beads coated with the indicated proteins. Data for the free microtubule ends and Ndc80+CENP-E coupled ends are from (*37*), plotted for direct comparison with CLASP2α and EB1. Columns (Mean ± SEM) are calculated based on N≥3 independent experiments and n examined microtubules (noted at each column) *p<0.05, ** p<0.001(Mann-Whitney u test), see Data source file for more details. C. Microtubule length vs. time (top) and the corresponding trajectory (bottom) of the distal end of a Hylite647-labeled microtubule segment relative to the CENP-E/Ndc80-containing bead (green circle, not to scale). This CENP-E/Ndc80-coupled end underwent three cycles of microtubule growth and shortening, while moving back and forth along a relatively straight path. D. Same as in panel C but for a typical bead coated with CLASP2α, which couples growing but not shortening microtubule tips (see Supplementary Fig. 9 for more examples). Coupled microtubule end grows persistently with small pauses, and pivots occasionally around the bead. E. Standard deviations of the pivoting angles around microtubule-bead attachment site for coatings with the indicated proteins. Box: first, second, and third quartile of the data set; whiskers: SD. Total number n of microtubule trajectories is indicated under each box. * p < 0.01 (Mann-Whitney u-test). F. Schematic of incomplete microtubule wall at the bead-coupled microtubule plus-end, which leads to softening of the bead-microtubule linkage and promotes visible microtubule pivoting.

Incorporation of tubulin dimers at microtubule plus-ends tethered by CENP-E and CLASP2α proceeded with occasional pausing and at a slightly reduced rate relative to the bead-free microtubules and the plus ends tethered by the beads containing CENP-E and either Ndc80 or EB1 (Fig. 8B). The catastrophe frequency of the CLASP2α-tethered ends was noticeably lower, indicating that the bead-conjugated CLASP2α is active and exerts stabilizing effects at the tethered microtubule tip (*5, 24, 37*). Unlike in traditional in vitro assays with free microtubule ends and soluble proteins, tethering dynamic microtubule plus-ends at the coverslip-immobilized beads has the benefit of providing insights into the mechanics of the dynamic tip structure. Indeed, thermally-induced pivoting of elongating microtubules around the attachment site is a sensitive gauge of the tip’s integrity because of a differing rigidity of different protofilament structures: individual protofilaments and their incomplete assemblies are much softer than a complete tubulin cylinder (*51*). We reasoned that if CLASP2α binding sites are at some distance from the tip, CLASP2α activity should have no effect on the rigidity of the bead-tip coupling, and the pivoting should be similar to that seen with other proteins that bind all along microtubule wall (Ndc80) or near the tip (EB). However, if the target for CLASP2α is at the tip’s extremity and the bound CLASP2α blocks tubulin addition to that site, microtubule tip integrity may become compromised.

We tracked the position of the distal end of microtubule seeds, which were moving away from the coupled beads owing to the tubulin incorporation at the bead-tethered ends (Fig. 8A). End-coupling via the CENP-E and the Ndc80 complex, which binds to lateral sites on the microtubule wall (*52, 53*), supported microtubule elongation along straight lines (Fig. 8C and Supplementary Fig. 9), implying a consistently rigid structure at the coupled end. A similar behavior was seen with EB1, which maintained the same tip structure while inducing frequent catastrophes (Supplementary Fig. 10), a property characteristic of soluble EB1 (*54*). In contrast, CLASP2α-coupled polymerization induced irregular pivoting around the microtubule attachment site (Fig. 8D), suggesting occasional softening of the bead-attached microtubule tip. Large deviations had variable durations and were seen only sporadically, indicating stochastic generation of the CLASP2 binding sites at the growing microtubule tip (Video 7). Such angular deviations were less frequent with Ndc80 (Video 8) and EB1 containing beads (Fig. 8E), suggesting that this phenomenon is specific to CLASP2α. We concluded that CLASP2α activity at the growing microtubule tip results in the occasional loss of complete protofilament structure. A sporadic loss of tip integrity is consistent with the idea that CLASP2α form high-affinity bonds with terminal GDP-tubulins and temporarily block assembly at one or several tubulin protofilaments (Fig. 8F).

## Discussion

Understanding the molecular mechanisms that regulate microtubule dynamics is both important and challenging. Great progress has been achieved using in vitro assays that examine microtubule dynamics in the presence of soluble forms of microtubule regulators. CLASPs, however, often function in cells as multimolecular ensembles concentrated near the microtubule tips, such as seen at mitotic kinetochores. By leveraging DNA origami technology and bead-tethering assays, we investigated the molecular activities of CLASP2α amplified by their clustering and without the interference from EB proteins, which are themselves potent regulators of microtubule structure and dynamics (*55*). Integrating these results with prior work by others (reviewed in (*6, 56*)) leads to a specific model for CLASP2-dependent stabilization of microtubule dynamics.

We propose that the TOG2 domain of CLASP2α has the ability to bind quickly and specifically to a GDP-tubulin dimer in the most extreme, terminal position of a polymerizing microtubule tip (Fig. 9A). Although dynamic microtubule ends elongate exclusively by incorporating GTP-tubulin dimers (reviewed in (*47*)), it is now well established that the structure and composition of the growing microtubule tip is highly complex and variable (reviewed in (*57*)). Terminal GDP-tubulins could arise at a growing microtubule tip via various mechanisms, e.g. through an occasional dissociation of the most terminal GTP-tubulin or accidental breakage of a vibrating single protofilament extension, as seen in computer models that account for protofilament mechanics under thermal forces (*58*). Although the microtubule tip has considerable stability and can tolerate several terminal GDP-tubulins (*58*), such “rogue” dimers are undoubtedly destabilizing and catastrophe-promoting because of the tendency of GDP-tubulins to disassemble, and also because they represent a poor site for incorporating new GTP-tubulins. A natural repair pathway, however, is available, whereby the GDP at the exposed E-site of terminal β-tubulin is replaced by GTP from a soluble pool (*48*). We propose that TOG2 of CLASP2α facilitates this exchange, thereby preventing tip erosion and indirectly promoting tubulin assembly (Fig. 9A). Justifications and possible implications of this model are discussed below.

**Figure 9.**
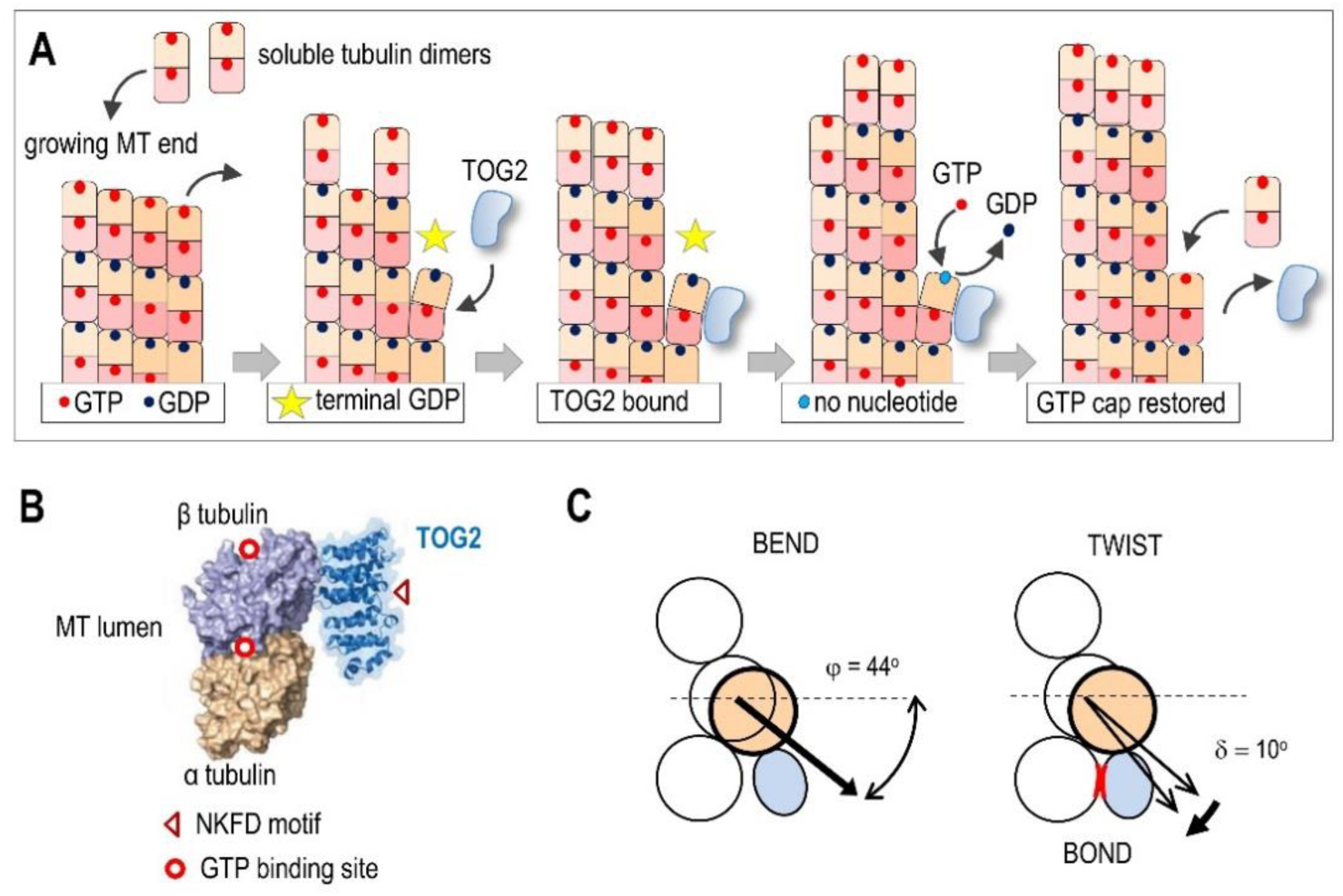
Molecular model for CLASP2α function at polymerizing microtubule plus-end. **A.** Two-dimensional schematic of a growing microtubule end illustrating the proposed cycle of TOG2 activity. TOG2 binds to terminal GDP-tubulin, which is shown to result from a stochastic dissociation of a distal GTP-tubulin. Bound TOG2 stabilizes this GDP-dimer against dissociation, while generating a one-protofilament nano-scale indentation at the tip. Spontaneous GTP exchange at the exposed E-site releases TOG2 and the “repaired” protofilament resumes its elongation. For simplicity, elongating protofilaments are depicted straight and GTP hydrolysis is depicted to occur immediately after the addition of distal GTP-tubulins, leading to a single-layer GTP cap. Two-dimensional model also fails to show correctly GDP-tubulin bending relative to the neighboring protofilaments, which is shown more accurately with a top-view in panel C. **B.** Structural model of human CLASP1 TOG2 domain complex with αβ-tubulin dimer (reproduced with modifications from (*27*)) showing approximate locations of GTP-bindings sites in tubulin monomers and NKFD motif in TOG2 domain. **C.** Schematic of three terminal tubulins (open circles) at the plus-end of microtubule cylinder viewed from that end. Left: yellow circle shows position of the middle β-tubulin monomer after it tilted away from the microtubule lumen at angle φ from the radial axis. The displacement shown corresponds to the intra-dimer bending angle of 18°, which should increase interaction surface with the TOG2 domain (blue) relative to the tubulin-TOG2 pair in panel B. Right: twisting motion of terminal GDP dimer may promote lateral contacts between the associated TOG2 domain and a dimer in the adjacent protofilament (red). Angles and displacements for this schematics were estimated based on (*27, 64*).

Structural work suggests that TOG2 binds along the tubulin dimer without hindering nucleotide exchange at the E-site of the associated β-tubulin (*27*) (Fig. 9B). In our assay, addition of soluble GTP caused CLASP2α dissociation from terminal tubulins with IC_50_ 320 nM, which is well below cytoplasmic GTP concentration. Thus, soluble GTP concentration should have no regulatory role for CLASP2α-tubulin interactions under the normal cellular environment. Instead, the proposed mechanism leverages the kinetics of GTP binding to terminal GDP-tubulin. For robust dynamic instability, the rate of GDP replacement within terminal tubulin dimers must be slower than the GDP-tubulin dissociation. Indeed, if the GDP-to-GTP exchange at the terminal tubulins were highly efficient, microtubules depolymerizing in the presence of GTP would quickly acquire terminal GTP-tubulins. The presence of polymerization-competent GTP-tubulins at the microtubule tip should lead to frequent pausing and even rescue, which is seen rarely with pure tubulin in vitro. This consideration suggests two possible mechanisms for TOG2-dependent microtubule stabilization via the GTP-dependent pathway. The parsimonious model that we currently favor is that TOG2 binding to terminal GDP-tubulin slows down its dissociation from the microtubule tip and buys time for spontaneous GDP-to-GTP exchange. Alternatively, TOG2 may have a catalytic, energy-consuming function in inducing polymerization-competent conformation of the terminal tubulin. In this more complex model, GTP binding or hydrolysis by TOG2 could be involved. We considered the latter model in view of the prior finding of a putative GTP-binding site within the only CLASP orthologue in *Drosophila* (*3*), however, our mutational analysis of human CLASP2α in vitro and in mitotic cells has ruled out that this site is involved. It is interesting in this context that the non-hydrolyzable GTP analogs have not been able to substitute for GTP in promoting TOG2 dissociation from microtubule tips in our assays, suggesting that putative GTP-ase activity of CLASP2α may require future investigation.

Our model, in which the TOG2 domain of CLASP family proteins stabilizes microtubule ends directly rather than by regulating GTP exchange or hydrolysis, is consistent with the prior finding that CLASPs do not affect the length of the GTP cap in microtubules growing in vitro (*38*). In the terminal-dimer binding mode to occasional rogue tubulins, CLASPs can exert stabilizing effects without significantly affecting the rates of microtubule growing, a behavior reported by multiple studies (*5*). Strong TOG2 binding to terminal GDP-tubulins also explains CLASPs’ inability to affect shortening rates and track with depolymerizing ends, since tip-tracking by soluble proteins requires weak and diffusive interactions (*59*). Prior mutational analysis demonstrated that conserved tubulin-binding residues in TOG2 domain abrogate both anti-catastrophe and rescue activities (*25*), suggesting that they are mediated by similar molecular mechanisms. Indeed, simultaneous binding of TOG2 molecules to several terminal tubulins could trigger the rescue of the depolymerizing end. Consistent with this proposal, Aher et al., observed brief CLASP2 binding events on shrinking microtubule ends immediately before rescue (*24*). Thus, the terminal binding model provides molecular insights into previous observations of CLASPs activity.

The proposed mechanism for CLASP activity also offers an appealing explanation for the apparent paradox that CLASP proteins are required for tubulin incorporation at the kinetochore-tethered ends, while at the same time they show no strong effect on the rate of tubulin addition at the freely-elongating microtubule ends. Indeed, at tethered microtubule ends, CLASP2α binding to terminal GDP-tubulins would assist capturing the ends of protofilaments that are prone to disassembly (Supplementary Fig. 11). Such unstable protofilaments are likely to arise frequently at the ends of kinetochore-tethered microtubules that experience persistent and variable force. A cluster of about 10 CLASP2α molecules in our assays provide hours-long binding to a microtubule end, ruling out the proposal that TOG2 causes dissociation of terminal tubulins from the tip. This collective, multivalent bond can withstand a pulling force up to 15 pN (Fig. 4); it may also stabilize the ends of microtubules under compression, as shown by (*24*). Thus, by forming molecular “nano-clasps” at the catastrophe-prone microtubule ends, clusters of CLASP2α localized at the kinetochores or in other intracellular locations could promote persistent elongation of tubulin protofilaments under adverse force conditions.

In our model, the target of TOG2 binding at the dynamic microtubule tip is any terminal GDP-tubulin, which is polymerization incompetent and does not support the incorporation of GTP-tubulin. A prediction of this model is that at the growing microtubule ends, TOG2 binding should correlate with the presence of protofilaments with stalled polymerization, giving rise to incomplete microtubule cylinders with altered mechanical properties. Consistent with this expectation, we observed occasional loss of integrity of microtubule tips polymerizing while tethered at the bead surface with CLASP2α and the plus-end-directed kinesin (Fig. 8). We surmise that such events are transient because after GTP is acquired at the stalled protofilament tip and TOG2 is released, the slightly lagging protofilament catches up quickly to restore typical microtubule end configuration (*60*). Also consistent with our model is a report of highly elongated protofilament protrusions with CLASP2α localizing at their base. Such unusual structures were observed in the comprehensive study by Aher and colleagues, who used a combination of soluble CLASP and EB proteins (*24*). This study concluded that CLASP2α protects growing microtubules against catastrophe when significant numbers of lagging protofilaments are present, in tune with our model. Microtubules growing in the absence of EB are likely to form protrusions at the nano-scale, escaping their visual detection but revealed via sensitive mechanical tests. The presence of EB may exaggerate this phenomenon, owing to EB activities at the microtubule tip, such as acceleration of both GTP-tubulin addition and its maturation into catastrophe-prone form (*54, 61–63*). Such dual activity would likely increase the occurrence of terminal GDP-tubulins with stalled TOG2-bound protofilaments, while at the same time promoting the incorporation of GTP-tubulin dimers at neighboring protofilaments. Similar effects likely explain why in the presence of EB, TOG2-S localizes behind the microtubule tip, which was defined by the most protruding protofilaments (*24*). Such localization is not incompatible with our model and it suggests that the TOG2 domain is actively engaged in “repairing” rogue GDP-dimers, which form frequently in the presence of EB. We also note that in addition to the proposed binding mode for TOG2, CLASPs may recognize some yet-unidentified feature of the GTP-cap or bind to curved protofilaments via other CLASP domains, such as TOG3 or SxIP-containing segment.

Our work highlights several interesting and still unresolved questions about the mechanism of TOG2-dependent stabilization of microtubules. Among the most enduring puzzles is the unusually bent shape of the TOG2 domain, which has been interpreted to prevent it from forming an extended binding interface with GDP-tubulin (*26, 28, 56*), in apparent contradiction to our model. However, a frequently overlooked property of tubulin dimers is their ability to undergo significant bending and twisting under thermal forces, as revealed by molecular dynamic modeling (*64*). The intra-dimer bending occurs on a microsecond scale with the maximum bending angle sufficient to provide contact with the convex surface of the TOG2 domain. We speculate that the highly curved surface of TOG2 domain serves to discriminate between terminal GDP- vs. GTP-tubulin dimers. Binding to the latter would lead to counterproductive interference with microtubule end dynamics, and it is likely avoided because GTP-tubulin is on average more straight than GDP-dimers (*65*). Thus, even though both dimers undergo frequent outward bending, only GDP-tubulin may exhibit such extreme conformations and become “captured” by TOG2. Modeling also suggests that GDP-tubulin that crowns the protofilament end bends outward with a significant twist, which may permit formation of a lateral bond between TOG2 and the neighboring protofilament (Figure 9C). We speculate that such three-step pathway – bend-twist-bond – underlies the stabilizing activity of TOG2 binding to terminal GDP-dimer. Analysis of such complex dynamic interactions will require theoretical modeling in combination with high resolution structural analysis and targeted mutations. We hope that our work and these ideas will guide and promote future research of the CLASP2 molecular mechanisms.

## ACKNOWLEDGEMENTS

Kinesin-1 proteins (K560) was generously provided by David Arginteanu and William Hancock (Penn State University). We thank Peter Relich and Melike Lakadamyali (University of Pennsylvania) for providing program for fluorescence intensity analysis. We thank Karla Miletic for assistance with image analysis; Emily Mattes, Po-Tao Chen, and Ekaterina Tarasovetc for help with protein purification, and Grishchuk lab members for discussions. We are also grateful to J.R. McIntosh (University of Colorado) for critical reading of the manuscript, and Rizal F. Hariadi (Arizona State University) for introducing us to DNA origami technology and providing help and reagents during initial stages of this project. Research reported in this publication was supported by the National Institute of General Medical Sciences of the National Institutes of Health under award number R35-GM141747 to E.L.G., R01 AI162260 and P50 AI150481 (via a collaboration development program of Pittsburgh Center for HIV Protein Interactions) to C.L., by the American Cancer Society grant RSG-14-018-01-CCG to E.L.G., by the European Research Council (ERC) consolidator grant CODECHECK, under the European Union’s Horizon 2020 research and innovation programme (grant agreement 681443), Fundação para a Ciência e a Tecnologia of Portugal (PTDC/MED-ONC/3479/2020), and a La Caixa Health Research Grant (LCF/PR/HR21/52410025) to H.M. F.I.A. acknowledges support from the Russian Science Foundation #21-45-00012. H.G. is a recipient of a PhD studentship from Fundação para a Ciência e a Tecnologia of Portugal (SFRH/BD/141066/2018).

## AUTHOR CONTRIBUTIONS

W.L. performed experiments using DNA origami technology; M.C. performed microtubule end conversion assays; V.D. and F.I.A. designed and performed experiments using optical trap; H.G. and H.M. designed and performed experiments using U2OS cells; Q.S. and C.L. designed and synthesized DNA origami scaffolds; A.M. constructed and purified CLASP2α truncated proteins, and carried out sequence analyses; E.L.G and W.L. designed research, analyzed data and wrote the paper with input from all authors.

## METHODS

### DNA origami construction and labeling

DNA origami ring structure was designed in caDNAno2 (caDNAno.org). The cylinder part was designed as in (*33*). Single strand DNA (ssDNA) handles extended from the 3’end of staple strands at positions indicated in Supplementary Fig. 1C and had the following sequences: 24 handles at the outer surface of the ring (AAATTATCTACCACAACTCAC ssDNA), 8 handles at the inner surface (CTTCACACCACACTCCATCTA ssDNA) and 4 handles on the top of the ring (CGGTTGTACTGTGACCGATTC ssDNA). The origami structures were assembled through 85 °C to 25 °C annealing gradient protocol for 36 hours in 1×TE buffer (10 mM Tris-HCl pH 8.0, 1 mM EDTA) supplemented with 10 mM MgCl_2_. Molecular weight of assembled DNA origami was 5 MDa. The assembled sample was purified by rate-zonal centrifugation through a 15–45% glycerol gradient in 1×TE supplemented with 10 mM MgCl_2_ in an SW 55 rotor (Beckman Coulter) at 303,800 g (RCF) at 4°C for 1 hr and fractions were collected and stored at -20°C. To carry out negative-stain electron microscopy, purified DNA origami sample was deposited onto glow-discharged 400 mesh formvar/carbon-coated copper grids (Electron Microscopy Sciences). Grids were then stained with 2% uranyl formate. Imaging was performed on a JEOL JEM-1400 Plus microscope operated at 80 kV with a bottom-mount 4k×3k CCD camera (Advanced Microscopy Technologies) (*33*).

Outer surface “anti-handle” oligos were generated using 5′-amino-labeled ssDNAs purchased from Integrated DNA Technologies (IDT, IA, USA). These oligos were labeled with benzyl-guanine (BG) by mixing together 4 µl of 5′-NH_2_-oligos at 2 mM, 8 µl of 200 mM HEPES buffer (pH=8.5) and 12 μl of BG-GLA-NHS (NEB, MA, USA, Cat. no. S9151S) prepared in DMSO at 20 mM. The mixture was incubated for 30 mins at 25 °C and ssDNA was precipitated by addition of 226 µl of 100% ethanol and 50 µl of solution containing 500 mM NH_4_Ac, 10 mM Mg(Ac)_2_ and 2 mM EDTA. The mixture was kept at -20 °C overnight, spun at 21,000 × *g* for 30 mins at 4 °C. Ethanol supernatant was aspirated; pellet was air dried, then resuspended in 200 µl H_2_O and stored in aliquots at -20 °C. Successful BG-modification for each preparation was confirmed by running a sample on a 20% polyacrylamide gel (Supplementary Fig. 1D).

For inner and top anti-handles, we used antisense 5′-Cy5-ssDNA and 5′-biotin-ssDNA, respectively (purchased from IDT). To attach anti-handles, DNA origami stock diluted to 5 nM with 1xTE buffer supplemented with 10 mM MgCl_2_ was mixed with 400 nM Cy5-oligos and 200 nM and biotin-oligos. The mixture was incubated for 2 h at 37 °C, diluted 10-times with 1xTE buffer supplemented with 10 mM MgCl_2_, aliquoted, snap frozen and stored at -80 °C. Concentration of DNA origami and ssDNA oligos was determined by measuring A260 with NanoDrop (Thermo Fisher Scientific, MA, USA).

### Construction of CLASP2α variants for in vitro studies

Truncated CLASP2 protein L-TOG2-S is analogous to the protein with the same name described in (Aher et al., 2018), but it was designed using human CLASP2α sequences (GenBank; NM_015097.3). It was constructed using synthesized DNA fragment with L-TOG2-S (from Lys261 to Ser793 amino acids, Genewitz) with codon usage optimized for expression in *E. coli*. Tags (6×His and GFP) and a linker containing SGGGGSGGGGSGGGG sequence were added at the 5’-end of the L-TOG2-S open reading frame, and subcloned between SpeI and HindIII restriction sites of the expression vector pRSETa, generating GFP-L-TOG2-S fusion protein. GFP-L-TOG2, containing sequence from Lys261 to Ser557 of CLASP2α, was generated by truncating the S-domain at the HindIII and PspXI restriction sites, followed by blunting with Klenow fragment and re-ligation. Site directed mutagenesis of GFP-L-TOG2-S was done at the Penn Genomics Analysis Core to generate GFP-L-TOG2-S-SKFS (containing N403S and D406S), and GFP-L-TOG2-S-NMFD with K404M.

### Protein purification for in vitro studies

Tubulin was purified from cow brains by thermal cycling and phosphocellulose (Poly Sciences, 19792-100) chromatography as in (*66*). Tubulin was labeled with Rhodamine or Hilyte647 as in (*67*). Human GFP-tagged Ndc80 ‘‘Broccoli’ was purified as in (*52*), GFP-tagged CENP-E kinesin motor domains as in (*68*); human EB1-GFP-6x His as in (*63*), SNAP-GBP as in (*35*). Drosophila Kinesin 1 (K560 construct) was purified and labeled with biotin (biotin-K560) as in (*69*). Purification of human full length GFP-CLASP2α was as in (*37*). Truncated CLASP2α constructs were expressed in *E. Coli* Arctic Express (DE3) cells (Agilent, CA, USA), grown in 2×YT broth (Thermo Fisher Scientific, MA, USA) to reach OD 0.9-1.0 and purified following protocol in (*24*) with modifications. Briefly, cultures were snap cooled on ice for 30 min, and protein expression was induced by addition of 0.5 mM 1-thio-β-galactopyranoside (LabScientific, NJ, USA) for 22-24 h at 10 °C with 240 rpm shaking. Cells were harvested by centrifugation at 5,000 × *g* for 30 min at 4 °C, then resuspended in lysis buffer: 50 mM HEPES, 500 mM NaCl, 10 mM imidazole, 5% glycerol, 2 mM β-mercaptoethanol, 0.5 mM AEBSF (GOLDBIO, MO, USA), and “Complete protease inhibitors cocktail” (Roche, Basel, Switzerland). Cells were lysed by sonication and a crude lysate was centrifuged at 22,000 rpm for 30 min (Ti50.2 rotor and optima XPN-80 ultracentrifuge, Beckman Coulter, CA, USA). Supernatant was loaded onto the HisTrap HP 1 ml Ni^2+^column operated with ÄKTA pure chromatography system (GE Healthcare, IL, USA). The column was washed with 20 column volumes of the lysis buffer without protein inhibitors, and protein was eluted via 10-250 mM imidazole gradient. Affinity purification was followed by size-exclusion chromatography using Superdex 200pg 16/600 column (GE Healthcare, IL, USA) equilibrated with SEC buffer (50 mM HEPES, 500 mM NaCl, 5% glycerol and 2 mM β-mercaptoethanol). Fractions containing target protein were pooled and concentrated using 10k MWCO centrifugal filters (Amicon, MA, USA). Protein was aliquoted with final 25 % glycerol, snap frozen and stored at -80 °C.

Prior to each experiment, a frozen protein aliquot was diluted 2-10 times in PBS (NaCl 137 mM, KCl 2.7 mM, Na_2_HPO_4_ 10 mM, KH_2_PO_4_ 1.8 mM, pH 7.4) and clarified by ultracentrifugation (TLA100 rotor, Beckman Coulter) at 27,904 × *g* for 15 mins at 4 °C. Concentration of GFP-tagged protein in the supernatant was determined by measuring GFP intensity by fluorescence microscopy and comparing it to a “standard” GFP-labeled protein whose concentration was determined by spectrometry, as described in (*50*).

### Fluorescence microscopy assays

In vitro assays using fluorescence imaging with no force application were carried out with an inverted Eclipse Ti-E Nikon microscope equipped with 1.49 NA 100× oil objective, Perfect Focus system and Andor iXon3 CCD camera (Cambridge Scientific, MA, USA), as in (*68*). Three diode lasers (488 nm, 561 nm and 640 nm from Coherent, CA, USA) were used as a light source with a C-TIRF Quad cube (Chroma, VT, USA). The multi-color total internal reflection fluorescence (TIRF) was implemented by rapidly switching between different laser wavelengths with an acousto-optical tunable filter and corresponding emission filter wheels. Under these conditions, the microscope produced 512 × 512-pixel images with 0.14 μm pixel^−1^ resolution in both the *x* and *y* directions. Illumination was carried out with a partially opened iris diaphragm to limit laser illumination only by the field of view during each exposure. Each imaging channel was recorded with 300 ms exposure time unless otherwise specified. All experiments were carried out at 32 °C by heating the objective with an objective heater (Bioptechs, PA, USA).

### Preparation of flow chambers with the coverslip-immobilized DNA origami

Coverslips (22 × 22 mm) were plasma cleaned and silanized with Repel-Silane ES (GE Healthcare Life Sciences, 17-1332-01), as in (*70*). To form a flow chamber, a silanized coverslip was attached to a regular glass slide using 4 strips of the double-sided tape (Scotch, MN, USA) as spacers, which formed 3 independent parallel flow channels with ∼10 µl volume each. Body-double fast silicone rubber (Smooth-On, PA, USA, Cat. no.83741) was painted on glass slide to create inlet and outlet grooves that prevented solution mixing between the channels. Solutions were exchanged by placing assembled chambers on a gently sloped surface, adding 15 µl at the inlet of each channel one by one and letting solutions to flow by gravity. Each incubation was for 10 min followed by washing with 50 µl of PBS-BSA buffer (PBS supplemented with 4 mg/ml BSA and 2 mM dithiothreitol (DTT)), unless specified otherwise. To immobilize DNA origami, we first incubated chambers with 22.5 µM biotinylated bovine serum albumin (biotin-BSA) (Sigma-Aldrich, MO, USA, Cat. no. A8549), then with 25 µM neutravidin (Thermo Fisher Scientific, MA, USA, Cat. no. 31000), followed by 5 pM of biotin- and Cy5-labeled DNA origami in TE buffer supplemented with 10 mM MgCl_2_; the surfaces were then blocked with 1% Pluronic F127 (Sigma-Aldrich, Cat. no. T2443).

To conjugate GFP-tagged proteins to coverslip-immobilized DNA origami, SNAP-GBP (5 μM) was incubated with BG-oligos (1 μM) in an Eppendorf tube for 1 h at 25 °C, diluted to 200 nM in Mg-BRB80-casein buffer containing 80 mM PIPES pH 6.9, 4 mM MgCl_2_, 1 mM EGTA, 0.5 mg/ml casein (Sigma-Aldrich, Cat. no. C5890), 4 mg/ml BSA and 2 mM DTT. The mixture was added to each channel, incubated for 10 min and washed with Mg-BRB80-casein. GFP-tagged protein (10 nM) in Mg-BRB80-casein buffer was incubated for 10 min, the chamber was washed with Imaging buffer (Mg-BRB80-casein buffer supplemented with up to 10 mM DTT, 6 mg/ml glucose (Sigma-Aldrich, Cat. no. g8270), 20 μg/ml catalase (Sigma-Aldrich, Cat. no. C40) and 0.1 mg/ml glucose oxidase (Sigma-Aldrich, Cat. no. G2133). The chamber was placed on the prewarmed microscope stage and imaged in GFP channel and Cy5 channels to collect images of 10-25 fields separated by at least 100 µm to avoid pre-bleaching. Typical settings for imaging of functionalized DNA origami in GFP and Cy5 channels were: 10 % intensity of 488 nm laser and 30% intensity of 640 nm laser. The settings for Andor iXon3 camera for both fluorophores were: gain 5.0, EM gain 300, 1 MHz readout speed, 300 ms exposure time.

### Determining number of GFP-containing protein molecules conjugated to DNA origami

Conjugation efficiency of GFP-tagged protein to DNA origami was calculated as a ratio of the total GFP intensity of DNA origami and the intensity of single GFP-tagged molecule. For semi-automatic analysis of total GFP intensity of DNA origami we used custom-written MATLAB program, which identified coordinates of local maximal pixel intensity in the Cy5 channel and then determined integrated fluorescence intensity of the square regions (7 × 7 pixel) centered on these coordinates in the GFP channel. This procedure eliminated selection bias and avoided contaminating our data sets with intensities of occasional protein aggregates.

Intensity of single GFP-labeled molecules was determined in the same experimental chambers by quantifying intensity of dim dots that did not colocalize with DNA origami. MATLAB program twobox based on DIPimage tool box (http://www.diplib.org/dipimage) (kindly provided by Dr. Peter Relich and M. Lakadamyali) selected local maximal pixel intensity in the GFP channel and collected integrated intensity I_7_ and I_9_ within two square regions centered on each selected dot: 7 × 7 pixels inner box and 9 × 9 pixels outer box. Data were collected in both GFP and Cy5 channels, and only GFP dots that lacked Cy5 brightness were used. Because of the selected regions geometry, background intensity in the GFP channel corresponds to average pixel intensity of the space between inner and outer boxes multiplied by the number of pixels in the inner box. Thus, integrated GFP intensity of each dot, *I_dot_*, can be calculated using the following equation:

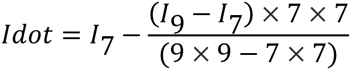

Intensity distribution for single molecules was then generated, and negative values (arising from image noise) were removed. Intensity of single GFP-labeled molecules was then calculated as a peak value of Gaussian fitting.

Because some proteins may cluster or aggregate, we additionally determined single molecule intensity using photobleaching, as in (*70*). Briefly, dim GFP dots that attached non-specifically in the DNA origami chambers were imaged for 2 min via stream acquisition at the rate of 300 ms per frame. Using Metamorph software (Molecular Devices, CA, USA), each clearly discernable dot was surrounded with the circle 8 pixels in diameter, while avoiding dots that were less than 4 pixels apart. After subtracting the background intensity, the intensity histogram for bleaching curves was fitted with equidistant Gaussian function, leading to a highly similar estimate of single molecule intensity as with the approach that used only initial brightness of GFP dots, rather than the entire photobleaching curve. The intensities of single molecules of Ndc80 Broccoli-GFP and GFP-CLASP2α determined with these methods were similar (Supplementary Fig. 2E), consistent with the predominantly monomeric form for these proteins at nanomolar concentrations in our assays (*24, 37, 71*). Also, the labeling efficiency we report (∼44%) is typical of DNA-origami directed protein assemblies, considering the handle incorporation rate, SNAP-to-BG reaction yield, and GFP-GBP binding affinity.

### Assays with stabilized microtubules and DNA origami-conjugated protein clusters

Taxol-stabilized microtubules were prepared as described in (*50*) using unlabeled tubulin (10 mg/ml) and HiLyte647-labeled tubulin (degree of labeling 0.18; 8 mg/ml) in a 5:2 ratio, rhodamine-labeled tubulin (degree of labeling 0.9; 6 mg/ml) in a 10:1 ratio, and 1 mM Mg-GTP (Sigma-Aldrich, Cat. no. G8877). GMPCPP-stabilized microtubules were prepared analogously but with 1 mM GMPCPP (Jena Bioscience, Jena, Germany, Cat. no. NU405L). After polymerization, microtubules were diluted 10 times with Mg-BRB80 buffer and spun at 16,000 × *g* at room temperature for 15 min. The pellet was resuspended, and the spinning procedure was repeated one more time to remove soluble tubulin. Microtubules were kept in Mg-BRB80 buffer supplemented with 10 µM taxol (Sigma-Aldrich, Cat. no. T7402) or 1 mM GMPCPP at room temperature.

To assay microtubule binding to the origami-based protein clusters, taxol-stabilized microtubules were diluted 1:10 in imaging buffer supplemented with 10 µM taxol and flowed into the chamber with functionalized DNA origami and incubated for 30 min. Images in Cy5 and GFP channels were captured to define position of origami-based CLASP2α clusters. Time-lapse video of microtubules was taken in rhodamine channel for 30 - 360 min, as indicated for each experiment, at 1 frame per 30 s (exposure 300 ms). The following imaging conditions were used: 1% intensity of 561 nm laser and camera settings gain 5.0x, EM gain 300, 1 MHz readout speed, 300 msec exposure time. Number of origami-bound microtubules was determined as a function of time and results were plotted as survival probability using the Kaplan–Meier algorithm implemented with Origin software (OriginLab, MA, USA). Lifetime of the tethered microtubules in the presence of different nucleotides was determined analogously.

To determine polarity of microtubules bound to the CLASP2α clusters, experiment was carried out analogously but using custom-made flow chambers with a syringe pump, enabling solution exchange at well-controlled flow rates and without chamber drying. After microtubules formed end-attachments to the origami-based CLASP2α clusters, biotinylated K560 kinesin (400 nM in PBS) was flowed into the chamber and incubated for 5 min to immobilize K560 on the neutravidin-coated coverslip surface. Then, Mg-ATP (10 μM, Sigma-Aldrich, Cat. no. A9187) in imaging buffer was flowed in and microtubule gliding was recorded using 561 nm laser. Kymographs of gliding microtubules were prepared with Metamorph and overlaid with static DNA origami images taken in GFP channel.

To investigate whether soluble tubulin can incorporate at the microtubule plus-end bound to clusters of CLASP2α, end-on attachments were formed between origami-bound CLASP2α and GMPCPP-containing rhodamine-labeled microtubule seeds via incubation for 5-10 min and a wash to remove unbound microtubules. Soluble Hilyte647-labeled tubulin (8 µM) was flowed at 25 μl/min in imaging buffer supplemented with 1 mM Mg-GTP. Images were taken alternating Hilyte647 (10% laser intensity) and rhodamine channels (1% laser intensity) for 300 s with 300 ms exposure time and 5 sec intervals, and the following camera settings: gain 5.0x, EM gain 300, 1 MHz readout speed.

### Rupture force measurements using optical trap

Our laser trap instrument has been described in (*72, 73*). Chambers with immobilized DNA origami were prepared and functionalized with full length CLASP2α as described above. Taxol-stabilized rhodamine-labeled microtubules were flowed in and incubated for 30 min at 32°C to promote formation of end-on attachments. After unbound microtubules were washed away with imaging buffer, beads coated with anti-tubulin antibodies were flowed in. Beads were prepared as in (*73*). Briefly, streptavidin-coated polystyrene beads (Spherotech, IL, USA, Cat. no. SPV-05-10) were incubated for 30 min at room temperature with 10 µg/ml of anti-tubulin antibodies (Biolegend, CA, USA, Cat. no. 801211) in PBS supplemented with 4mg/ml BSA and 2 mM DTT. Beads were washed by centrifuging at 4,500 × *g* for 7 min at 4 °C, incubated with 0.8 mM of b-PEG (Quanta BioDesign, OH, USA, Cat. no. 10776) to block bead surface and stored at 4 °C for no more than 2 months. Prior to each experiment 10 µl of beads were washed, resuspended in 10 µl of imaging buffer prior to introducing into the chamber; the chamber was sealed with Kwik-Cast Sealant (World Precision Instruments, KWIK-CAST) and placed on microscope stage prewarmed to 32 °C.

Tethered bead was found using Differential Interference Contrast imaging. To confirm that the bead is bound to a single microtubule, the bead was pulled gently with a low-power optical trap and the bead-bound microtubule was imaged briefly in rhodamine channel. Next, the bead was released from the trap and microtubule images were acquired in a Z-stack with 200 nm step. This procedure was necessary because most origami-bound microtubules were oriented perpendicularly to the coverslip plane, so it was not possible to capture microtubule image in one focal plane. Image of the coverslip surface was then acquired in GFP channel to confirm co-localization of the CLASP2α protein cluster and tethered microtubule end. Next, the tethered bead was recaptured in the optical trap at the appropriate depth to avoid exerting any forces on the bead. The piezo-stage (Physik Instrumente, Karlsruhe, Germany) was moved along *y*-axis at 20 nm/s for 1-2 min and bead coordinates were recorded with quadrant photo detector operated at 5 kHz. After the stage stopped moving, bead, microtubule and GFP-CLASP2α images were acquired again to determine the outcome.

To take into account that tethered bead moved in both *y* and *z* directions, total force acting on the bead was calculated as a square root of the sum of squared forces along these axes. Calibration of quadrant photo-detector was carried out as in (Barisic et al., 2015) using free-floating beads moved in *x* and *y* directions with acoustic-optical deflector and coverslip-immobilized beads moved along *z* axis with the piezo-stage. Trap’s stiffness along *y* and *z* axes measured using power spectrum method (Svoboda & Block, 1994) was 0.078 ± 0.003 and 0.029 ± 0.003 pN/nm, correspondingly.

### Assays with coverslip-bound stabilized microtubules

Anti-tubulin antibodies (Biolegend, CA, USA, Cat. no. 801211) were diluted 1:20 in PBS and flowed into microscopy chamber for 15 min at room temperature. Chamber was washed with Mg-BRB80 and coverslip surface was blocked by the addition of 1% Pluronic F-127 in Mg-BRB80 for 10 min. Chamber was washed thoroughly, taxol-stabilized microtubules labeled with Hilyte647 were added for 10 min, after which the unbound polymers were removed by washing. GFP-tagged protein of interest was flowed into the chamber and incubated for 10 min 32 °C. Chamber was sealed and moved to microscope stage prewarmed to 32 °C. Typically, we collected images for GFP and Hilyte647 wavelengths for 25 fields of view. Imaging of one microfluidics channel took 2-3 min, and all 3 channels were imaged one after another. Images of CLASP2α decoration were acquired using same settings as for other experiments with GFP-tagged CLASP proteins. Images of Ndc80 Broccoli-GFP were acquired analogously. At higher soluble concentration of Ndc80, EM gain was reduced 30-fold to avoid saturation.

Freshly prepared microtubules were used to minimize appearance of CLASP2α-containing foci along microtubule walls, presumably at the damage sites (*74*). On the day of experiment, CLASP2α protein aliquot was thawed, diluted to 100 nM with PBS buffer and ultra-centrifuged or 15 min at 27904 × g using Beckman TLA 100 rotor to remove protein aggregates; the supernatant was collected and kept on ice. GFP-protein concentration was determined in each chamber via GFP fluorescence (*50*).

Guanosine-containing nucleotides were prepared as Mg-containing salts. Briefly, powder of GTP, GDP (Sigma-Aldrich, Cat. no. G7127), GMPCPP (Jena Bioscience Cat. no. NU-1028), or GTPγS (Sigma-Aldrich Cat. no. G8634) was dissolved in 50 mM MgCl_2_ to prepare nucleotide stock solution at 50 mM (or 10 mM for GTPγS). Then, pH was adjusted to 7 with NaOH, as verified with pH test paper. The prepared stock solutions, as well as the commercial stock of GMCPPP (10 mM) were diluted to 100 μM in Mg-BRB80, aliquoted and stored at -80 °C. On the day of experiment, nucleotide stock was thawed and further diluted with imaging buffer to 1 µM. This concentration was chosen only ∼3-fold higher than the IC_50_ for GTP in order to reduce the effect from possible GTP contamination in these reagents. In assays examining effect of different GTP analogs on CLASP2α interactions with microtubules, nucleotide-containing buffer was added to microscopy chamber with immobilized microtubules and incubated for 30 min to allow ample time for nucleotide binding to tubulin. GFP-tagged full length or mutant CLASP2α protein was diluted to 1 nM with imaging buffer containing specified nucleotide and also incubated for 20 min to allow time for possible GTP binding prior to adding to microscopy chamber with immobilized microtubules.

To determine identity of the microtubule end decorated brightly with GFP-labeled CLASP2α, chambers prepared with silanized coverslip were incubated with biotin-BSA, then neutravidin, followed by incubation for 30 min with biotin-K560 (400 nM). Taxol-stabilized microtubules labeled with Hilyte647 were added in imaging buffer supplemented with 1 nM GFP-CLASP2α and 1 mM Mg-ATP. Gliding microtubules were imaged by alternating Hilyte647 (3% laser intensity) and GFP (10% laser intensity) channels for 120 seconds with stream acquisition (300 ms exposure for each channel) and using the following camera settings: gain 5.0x, EM gain 300, 1 MHz readout speed.

Cy3-GTP (Jena Bioscience, Cat. no. NU-820-CY3) was diluted to 100 nM in imaging buffer with or without 1 nM GFP-CLASP2α and incubated with coverslip-immobilized taxol-stabilized microtubules for 10 min. Images of Hilyte647-labeled microtubules and Cy3-GTP decoration were taken for 25 fields.

### Analysis of microtubule-end decoration

Images of coverslip-immobilized microtubules decorated with indicated GFP-tagged proteins or Cy3-GTP were analyzed using general procedures described in (*75*) with modifications. Briefly, only central quadrant of each image was analyzed to reduce any effects from uneven illumination. Images were examined visually to select non-overlapping microtubules that were longer than 2 µm and with both ends clearly visible. To avoid any bias, both microtubule ends were selected in Hilyte647 channel by placing 7 × 7 pixel boxes at both ends. A custom-made MATLAB program was then used to measure integrated fluorescence intensity within each box in the corresponding GFP (or Cy3) image. Background intensity in the same field of view was measured in three manually selected microtubule-free regions (7 × 7 pixel). Their average integrated intensity was subtracted from intensity measured on microtubules in the same field. At least 40 microtubules were analyzed for each experiment, and median intensity of the brightest microtubule end was determined. Apparent dissociation constant *K_d_* for GFP-CLASP2α was determined using one-site-binding model: *I* = *I_max_* × c/(*K_d_* + c), where *I* is intensity of microtubule end, *I_max_* is a plateau intensity and c denotes soluble GFP-CLASP2α concentration.

### Establishment of stable cell lines and RNAi treatment

The wild-type and SKFS (N403S, D406S) mRFP-CLASP2γ constructs were inserted between NdeI-NotI sites in a CSII-CMV-MCS vector (RIKEN BRC DNA BANK; RDB04377). The SKFS construct was generated from the wild-type construct by site-directed mutagenesis, using two overlap primers bearing the desired mutations. Lentiviral supernatants were produced in HEK293T cells using pMD2.G (Addgene, MA, USA, Cat. no. 12259), psPAX2 (Addgene, Cat. no. 12260) and the CSII-CMV-MLS plasmids, in the presence of Lipofectamine 2000 (Invitrogen, CA, USA) and Opti-MEM (Gibco, MD, USA).

A U2OS cell line stably expressing Photoactivatable (PA) GFP-α-tubulin (*76*) was transduced with the viral supernatants, in the presence of 6 μg/mL Polybrene (Sigma-Aldrich). The cell lines were cultured in DMEM with 10% FBS, supplemented with 10 µg/µL of antibiotic-antimycotic mixture (Gibco, MD, USA) and selected with Zeocine 1 µg/µL. Cell lines were maintained at 37 °C in a 5% CO2, humidified atmosphere. Cells expressing the different mRFP-CLASP2γ constructs were sorted using a FACS Aria II cell sorter (Becton Dickinson, NJ, USA). Briefly, cells were re-suspended in Basic Sorting Buffer (PBS 1X, Ca^2+^ and Mg^2+^ free; 5 mM EDTA; 25 mM HEPES; 2% FBS). Cells expressing red fluorescence were sorted into a tube containing DMEM with 10% FBS and then transferred to an appropriate growth flask. RNAi experiments were performed using Scrambled siRNA CUUCCUCUCUUUCUCUCCCUUGUGA, CLASP1α siRNA GCCAUUAUGCCAACUAUCU, or CLASP2α siRNA GUUCAGAAAGCCCUUGAUG with Lipofectamine RNAiMax (Invitrogen), as described in (*77*).

### Cell imaging and spindle measurements

All experiments were performed in the presence of MG132 (5 µM) added 30 mins prior to imaging to ensure that cells were in metaphase and prevent mitotic exit. To measure spindle length, cells were fixed, and microtubules were stained using mouse anti-α-tubulin clone B-512 at 1:1500 (Sigma Aldrich) and secondary antibodies Alexa Fluor anti-Mouse 488, diluted in PBS-Triton X-100 0.1% with 10% FBS. DNA staining was achieved by addition of 1 μg/ml 4’,6-diamidino-2-phenylindole (DAPI; Sigma-Aldrich).

Live-cell imaging was used to measure microtubule turnover rate. A thin strip of spindle microtubules was locally photoactivated in one half-spindle by pulsed near-UV-irradiation (405 nm laser; 500 μs exposure time). Fluorescence images (seven 1 µm-separated z-planes centered at the middle of the mitotic spindle) were captured every 15 s for 4.5 min with a 100x oil-immersion 1.4 N.A. plan-apochromatic objective. Under these conditions, the photoactivated region dissipated, yet the photoactivated molecules were retained within the cellular boundaries defining the cytoplasm. To calculate microtubule turnover, the sum intensity at each time point was fit to a double exponential curve A_1_×exp(-k_1_×t) + A_2_×exp(-k_2_×t) using Matlab (Mathworks), in which t is time. A1 represents the less stable population (typically interpreted as the fraction corresponding to non-kinetochore microtubules) and A2 represents the more stable population (typically interpreted as the fraction corresponding to kinetochore microtubules), with decay rates of k1 and k2, respectively. The half-life for each process was calculated as ln2/k for each microtubule population. Spindle poleward flux rates were determined in the same cells used for quantification of spindle MT half-life by determining the slope of the fluorescence signal over time (4.5 min), using a Matlab custom-written routine in LAPSO software (*78*) and using the slope of the movement of the metaphase plate as reference. Other details were as described in (*77*).

### Tethered-microtubule end assay with protein-coated beads

Experiments were carried out as in (*37, 50*). Motion of stable microtubule fragment away from the bead was scored if after addition of soluble tubulin there was a visible gap between the fluorescently labeled microtubule segment and the bead. The fraction of dynamic microtubule attachments was calculated as the ratio of number of microtubules that showed at least one away motion to the total number of microtubules bound to beads within a particular field. Velocities of the away and backward motions, their respective duration and the maximum lengths of polymerized microtubules for each dynamic cycle were estimated from kymographs. The total dynamic microtubule attachment time was calculated as time between the start and end of all dynamic motions, or the detachment of the seed from the bead, or the end of imaging. microtubule catastrophe frequency was calculated by counting the observed number of catastrophes and dividing it by the sum of the elongation time for all polymerizing microtubules. Two dimensional trajectories for motions of the leading microtubule ends were obtained using point tracking tool in Metamorph. Positions (*x,y* coordinates) of the bead-distant end of the labeled seed relative to the attachment site on the bead were used to calculate pivoting angles *θ = arc tan (y/x)* for each point, and the resulting histograms were approximated with Gaussian function. Whenever possible, data are presented as average of N independent trials, i.e. experiments carried out in separate microscopy chambers on different days. Small n denotes total number of observed events in all trials for each experimental condition.

## Abbreviations

BG: Benzylguanine
GFP: Green Fluorescence Protein
GBP: GFP binding protein
ssDNA: single stranded DNA
TIRF: Total Internal Reflection Fluorescence

## LEGENDS TO SUPPLEMENTARY FIGURES

**Supplementary Fig. 1.**
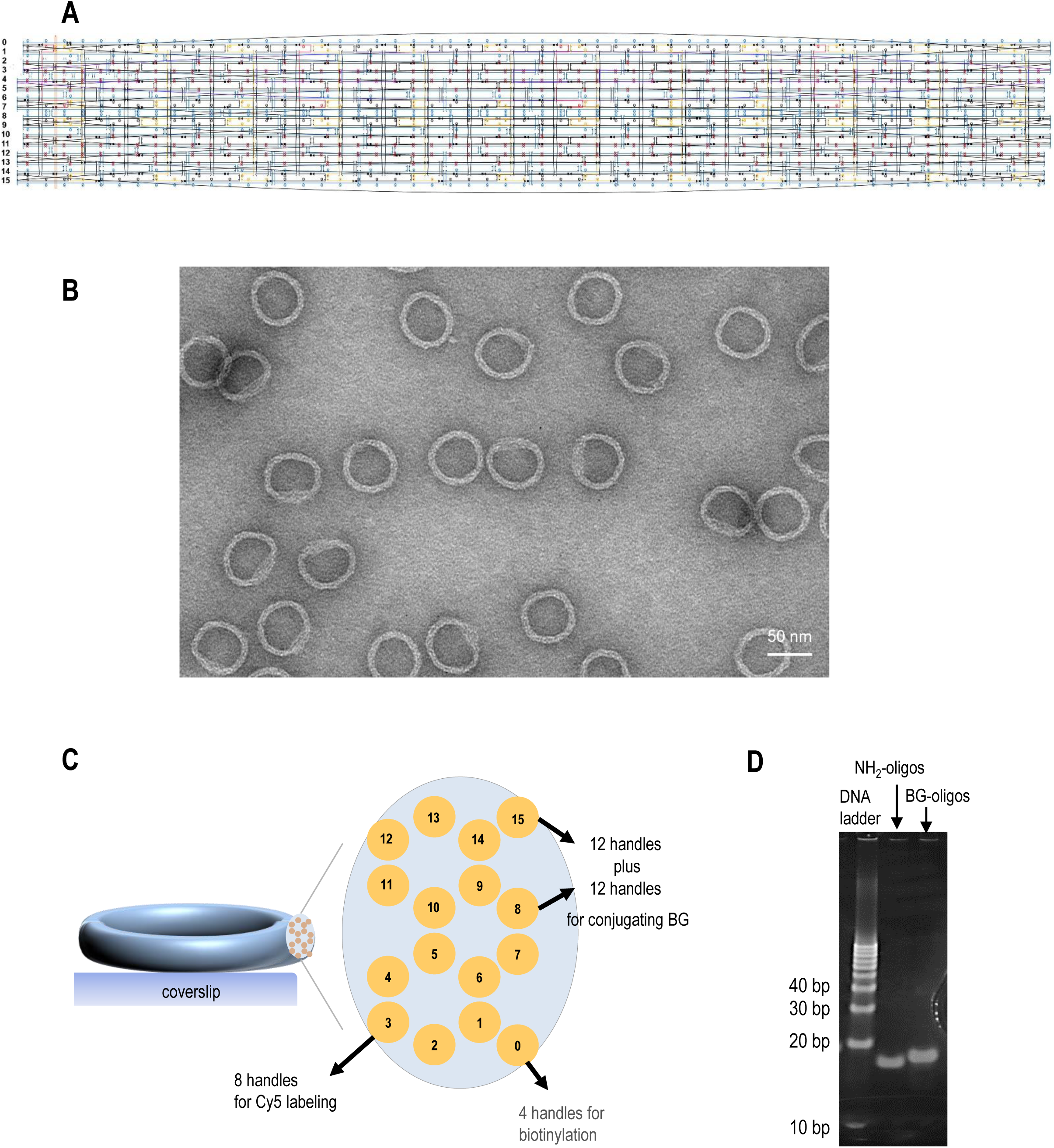
Location of DNA origami handles and BG-oligos preparation. **A.** DNA-origami designs rendered in caDNAno. The scaffold strand is in blue. Staples carrying 4 top handles, 8 inner handles, and 24 outer handles are shown in red, purple and orange, respectively. Handles (see Methods for sequences, not shown here for clarity) were extended from the 3’ end of staple strands. **B.** Negative-stain TEM image of the DNA-origami circles. Scale bar, 50 nm. **C.** Schematic shows a cross-section of DNA origami ring with numbered DNA strands (yellow circles). Handles are distributed evenly along the ring’s perimeter with total number of unique handles indicated. **D.** Image of a polyacrylamide gel stained with SYBR Gold shows NH_2_-oligos before and after reaction with BG-NHS. Successful labeling with BG is evident from the band shift to a higher molecular weight.

**Supplementary Fig. 2.**
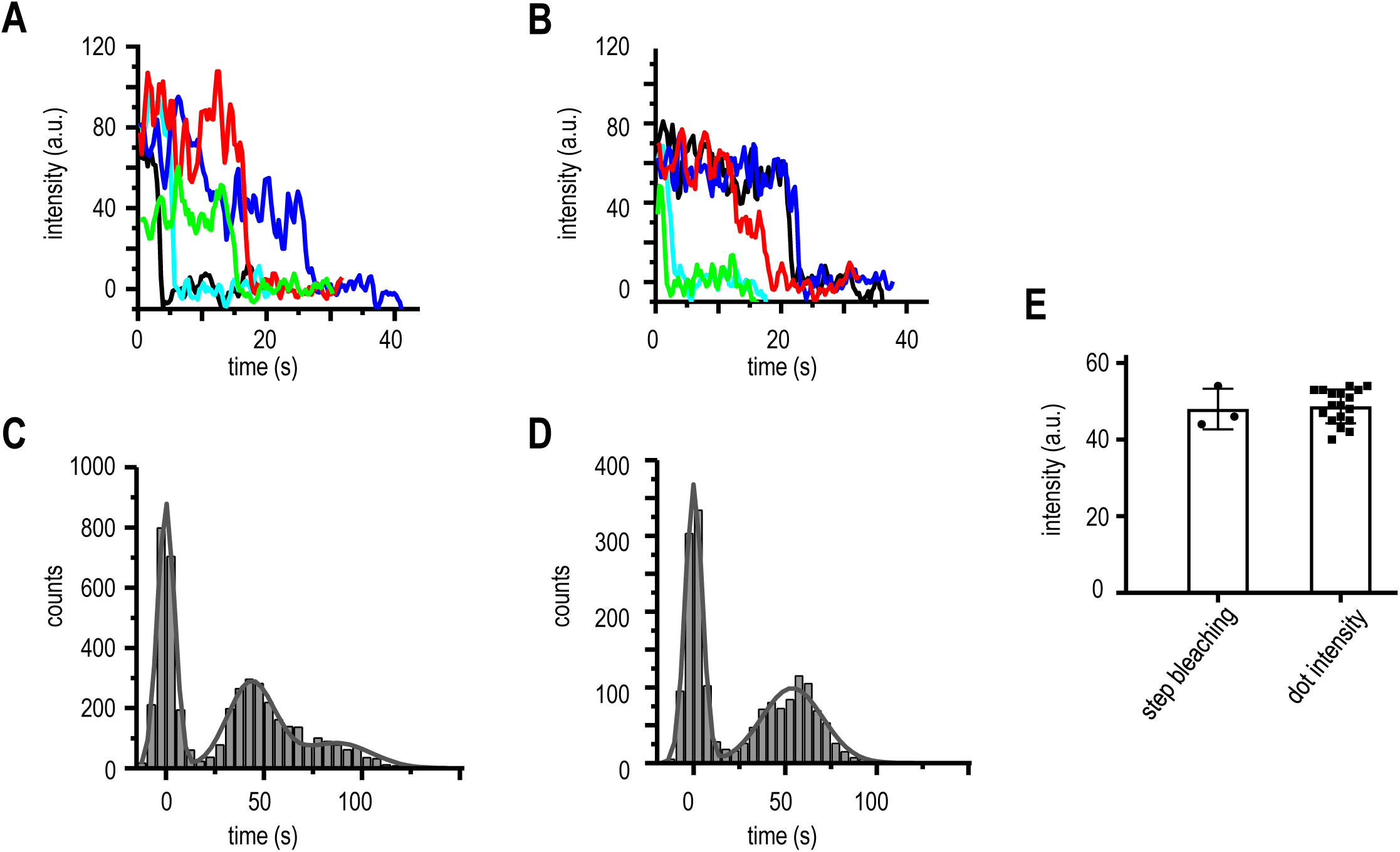
Single GFP intensity measured through photobleaching analysis. **A.** Representative photobleaching curves for non-specifically attached GFP-CLASP2α molecules. **B.** Same as in panel **A** but for Ndc80-GFP Broccoli. **C.** Distribution of intensities measured during photobleaching of GFP-CLASP2α molecules. Lines show equidistant Gaussian curves. Distance between the peaks corresponds to average single molecule fluorescence intensity. **D.** Same as in panel **C** but for Ndc80-GFP Broccoli. **E.** Average single molecule fluorescence intensity acquired via the photobleaching analysis, as in panels **A-D**, and by collecting initial brightness of non-specific dots on the coverslip during experiments with origami scaffolds. Each point on the graph corresponds to average intensity determined in an independent experiment. Data for non-specific dots brightness is the same as in Figure 2E but with no normalization, see Data source file for more details. Bars show mean ± SEM.

**Supplementary Fig. 3.**
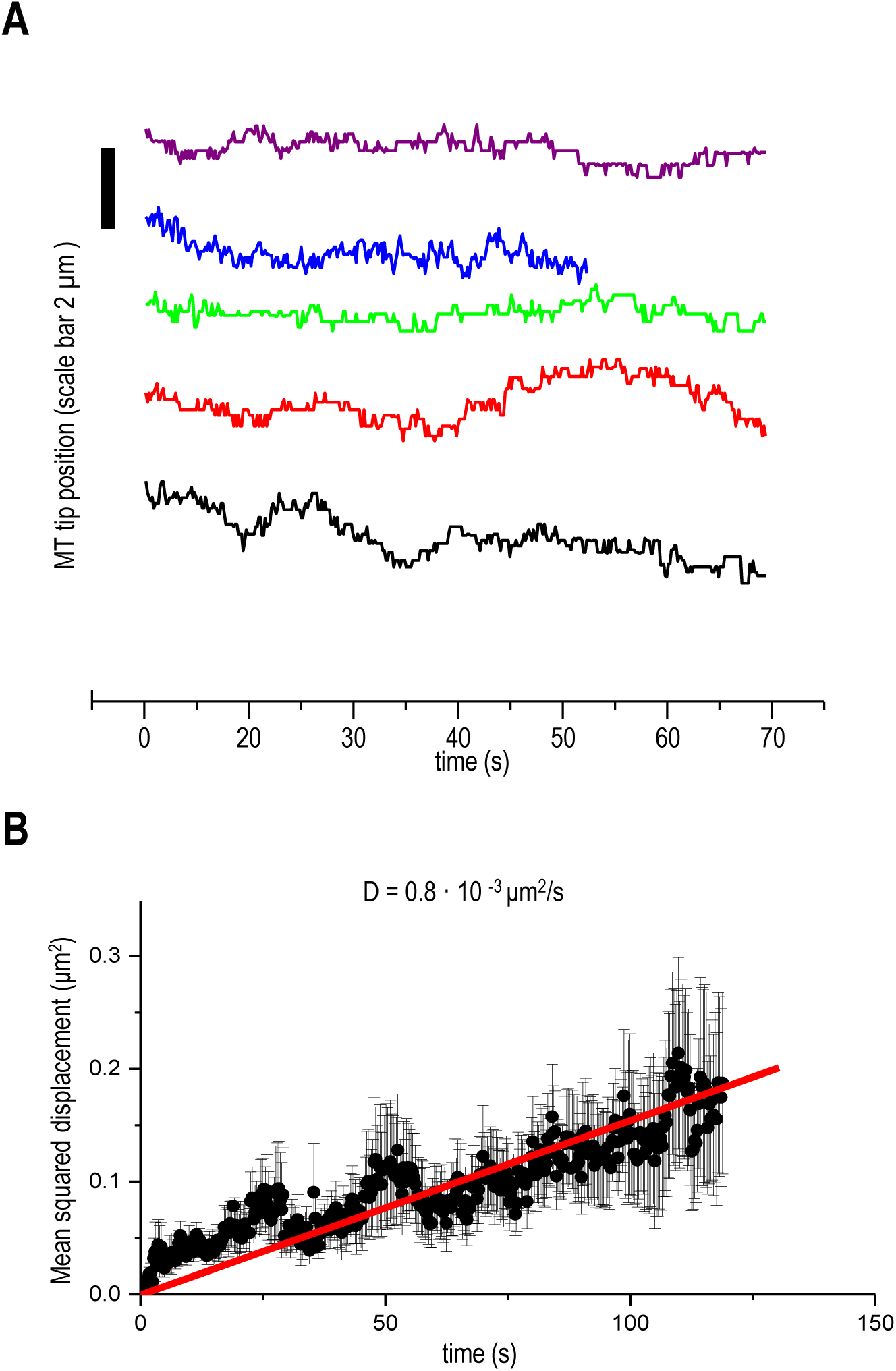
Origami-conjugated Ndc80 proteins support microtubule wall diffusion. **A.** Representative tracks for microtubules diffusing on DNA origami-based clusters of Ndc80-GFP Broccoli. **B.** Mean square displacement for 20 microtubules with a liner fit (red line) and calculated diffusion coefficient.

**Supplementary Fig. 4.**
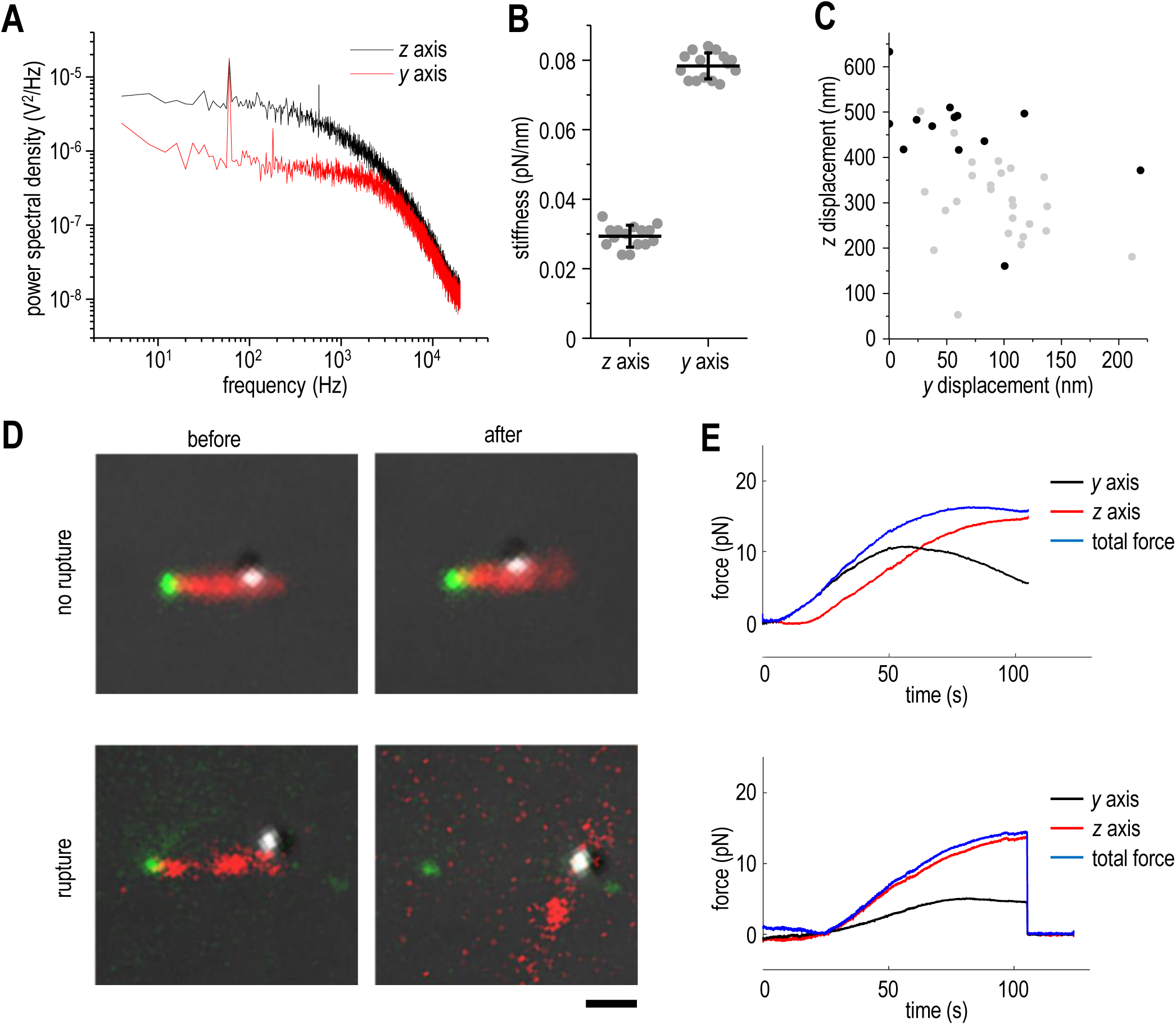
Rupture force experiments to examine the strength of CLASP2α attachment to microtubule plus-end. **A.** Lorentzian power spectrum for a 0.5 µm polystyrene bead in a static optical trap along (*z* axis) and perpendicular (*y* axis) to the laser beam. **B.** Stiffness of the optical trap along *z* and *y* axes. Mean ± SD, n = 15 beads, each point corresponds to individual bead. **C.** Bead displacement at maximal force for experiments that ended with rupture (black symbols) and no rupture (gray symbols). The displacements along *z*-axis are significant for many experiments, so there were taken into account when calculating total force acting on each bead. **D.** Representative images illustrating experimental outcomes. Left images show polystyrene bead (grey) bound to microtubule (red) which has its end tethered to a cluster with CLASP2α (green). Images on the right show same fields but after force application with top images representing a lack of rupture, and the bottom images showing a rupture between the cluster of CLASP2α and microtubule end. Bar is 1 μm. **E.** Force application curves for two example experiments show contributions of *z* and *y* components. Data were acquired at 5 kHz and smoothed by averaging with sliding window of 50 points. Top and bottom graphs represent experiments with no rupture and with a rupture, correspondingly. Total force was calculated as a square root of sum of squares of the *z* and *y* force components.

**Supplementary Fig. 5.**
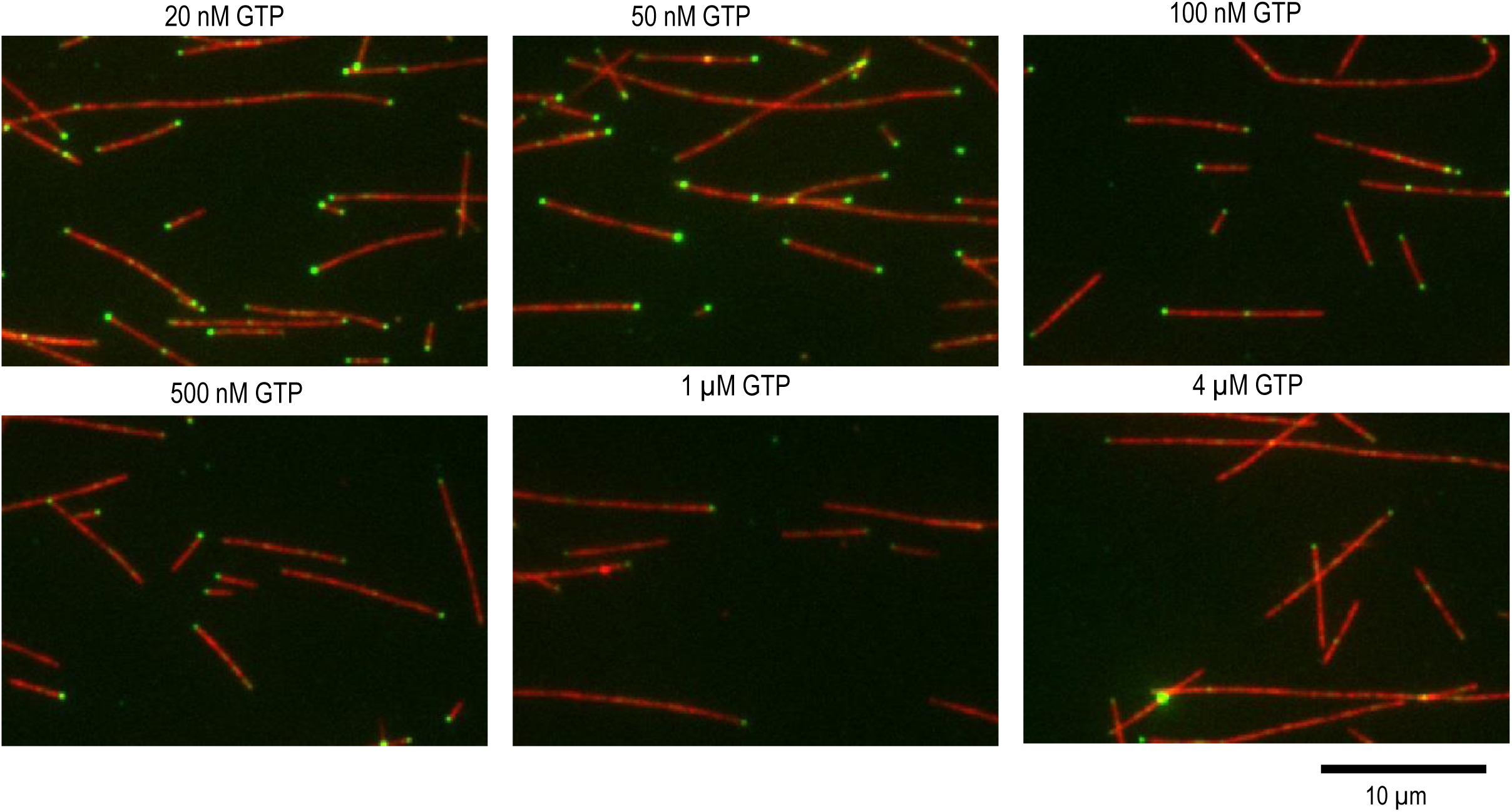
Microtubule decoration by GFP-CLASP2α in the presence of soluble GTP. Representative TIRF mages of taxol-stabilized microtubules (red) in the presence of 1±0.2 nM full length human GFP-CLASP2α (green) and indicated concentration of Mg-GTP.

**Supplementary Fig. 6.**
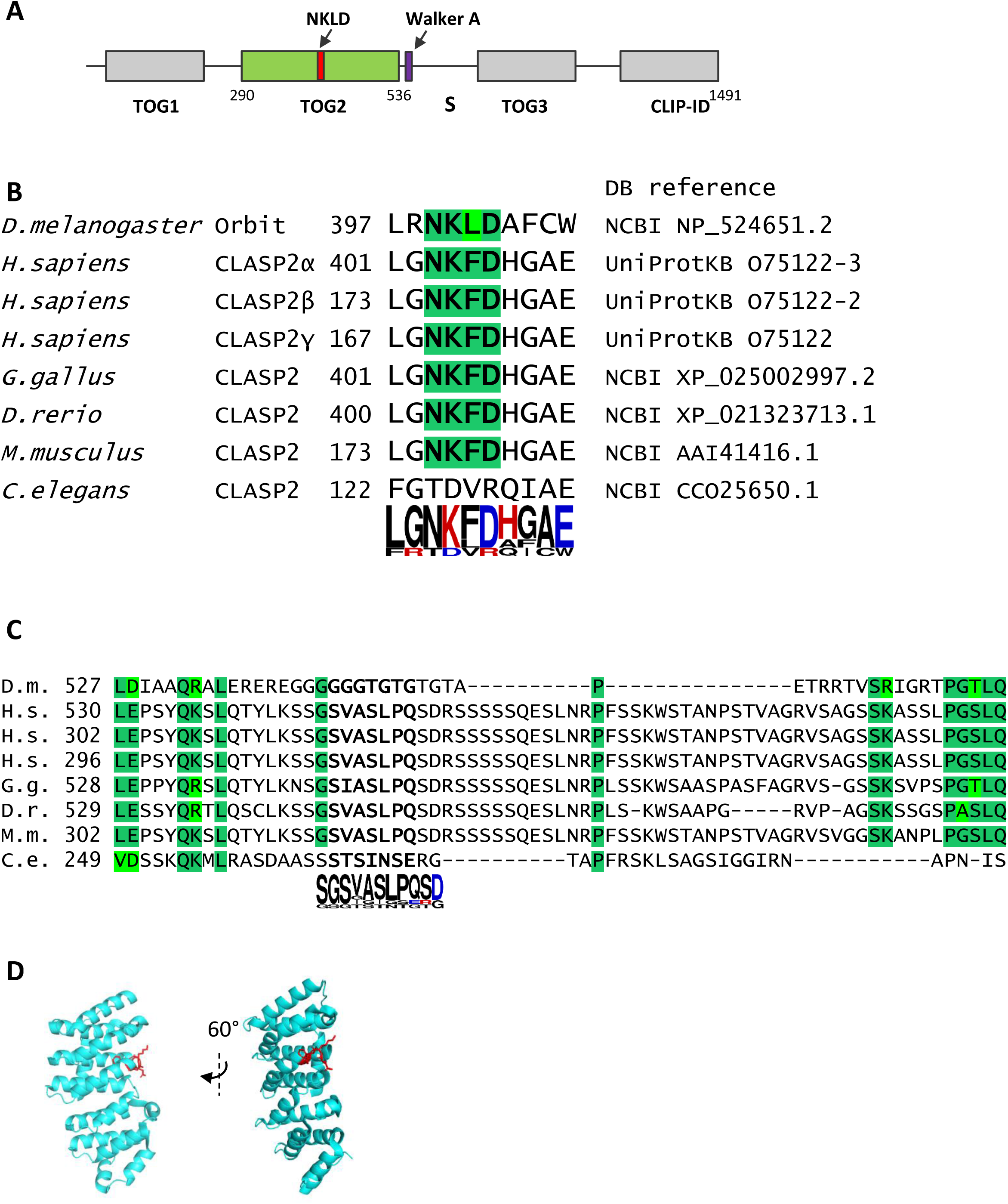
CLASP protein sequence and alignments. **A.** Domain structure of *Drosophila* Orbit/Mast (uniport sequence Q9NBD7)and location of canonical sequences for GTP binding, as identified in (3). **B.** Alignment of protein sequences for CLASP2 orthologs from different species at the sites with NKxD motif. Sequences were aligned using ClustalO; numbers at the start of sequences correspond to amino acids in original protein sequences. Most proteins contain NKDF sequence, with dark green color showing identical amino acids and light green showing conservative replacements. In a logo-representation at the bottom of alignments, size of the letter corresponds to the frequency of amino acid encounter. Color shows amino acid charge (blue for negative, and red for positive). **C.** Same as in panel B but showing protein alignment at the Walker A-like sequence. Protein sequence of *Drosophila* Orbit/Mast deviates significantly from sequences of other CLASP2 proteins and Walker A motif is lacking at this site. **D.** PDB structure (3WOY) of *H. sapiens* TOG2 with NKFD motif shown in red.

**Supplementary Fig. 7.**
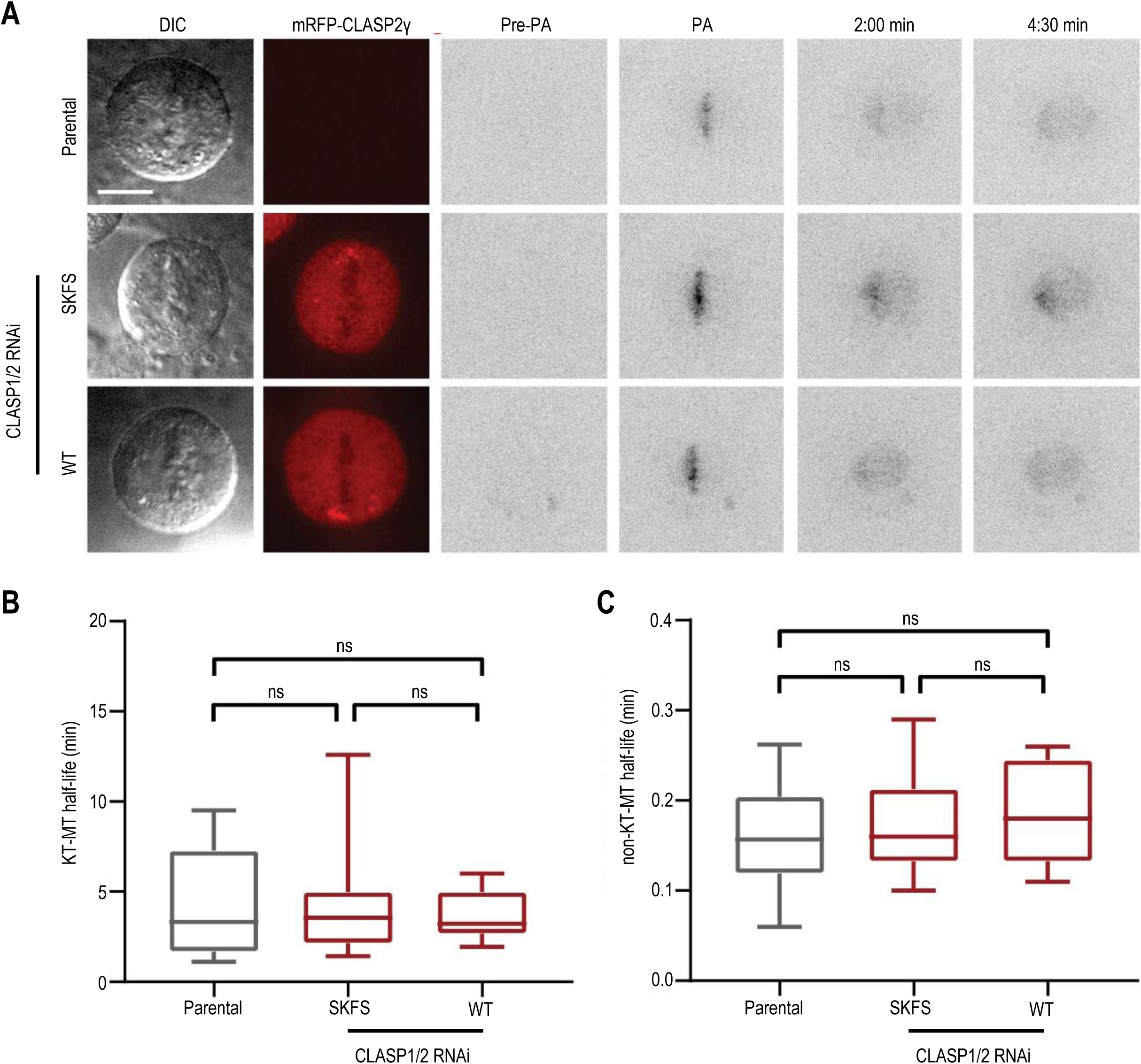
Lack of the strong effects from the SKFS mutation of CLASP2 in dividing cells. **A.** Photoactivation experiments in live U2OS parental PA-GFP-α-tubulin cells expressing the mRFP-CLASP2γ constructs after CLASPs RNAi; Panels represent DIC, the mRFP-CLASP2γ signal (red) and the PA-GFP-α-tubulin signal before photoactivation (Pre-PA), immediately after photoactivation (PA) and at 2 and 4.5 min after photoactivation. The PA-GFP-α-tubulin signal was inverted for better visualization. Scale bar is 5 µm. **B, C**. Quantification of half-life of spindle microtubules in the studied cell lines: kinetochores microtubules (KT-MT, panel B) and non-kinetochore microtubules (non KT-MT, panel C). See legend to Figure 6H for more details.

**Supplementary Fig. 8.**
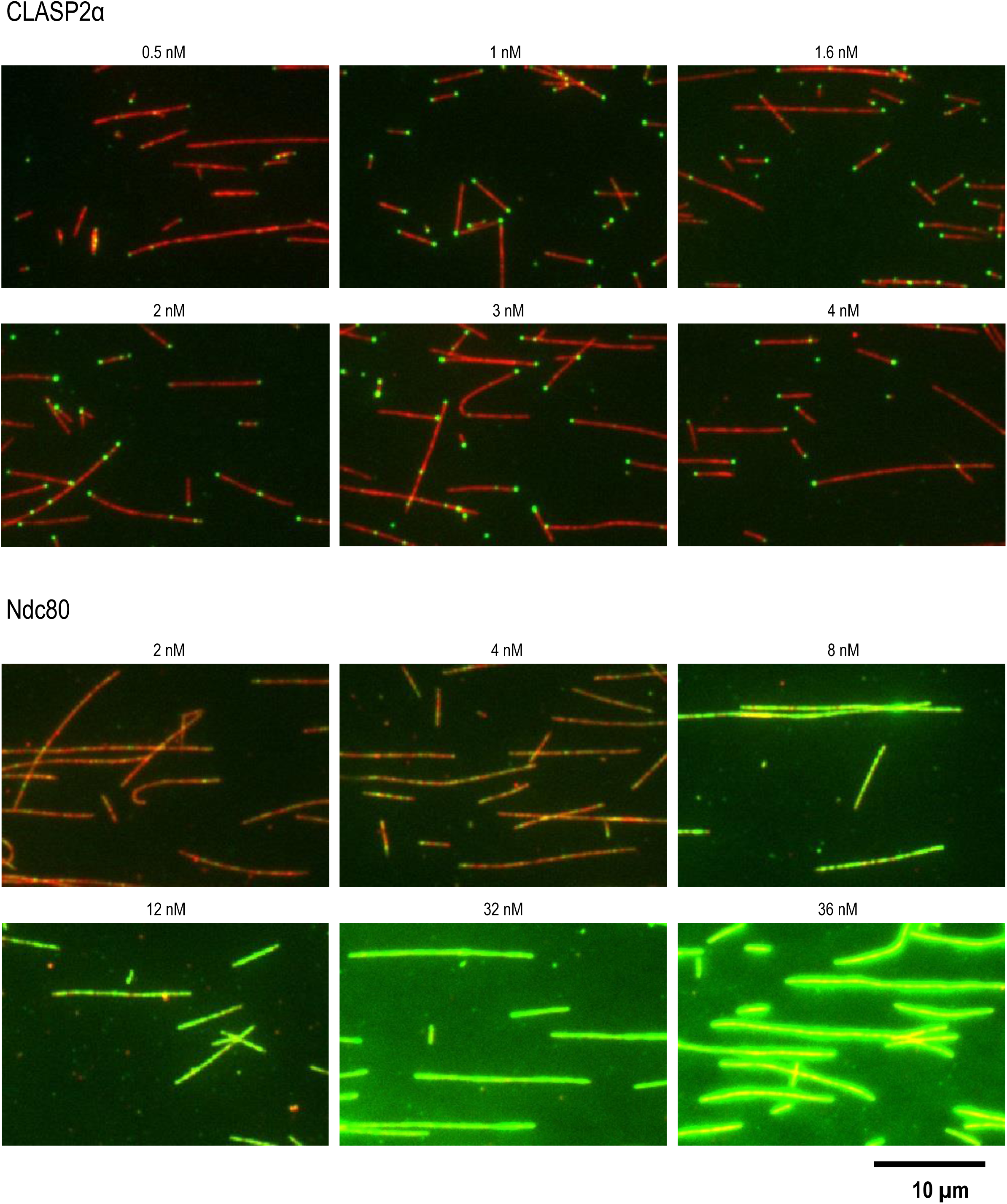
Microtubule decoration by GFP-tagged proteins. Representative TIRF images show taxol-stabilized microtubules (red) incubated in the presence of indicated concentrations of GFP-tagged full length human CLASP2α (top set of images) or human Ndc80 Broccoli (lower set of images).

**Supplementary Fig. 9.**
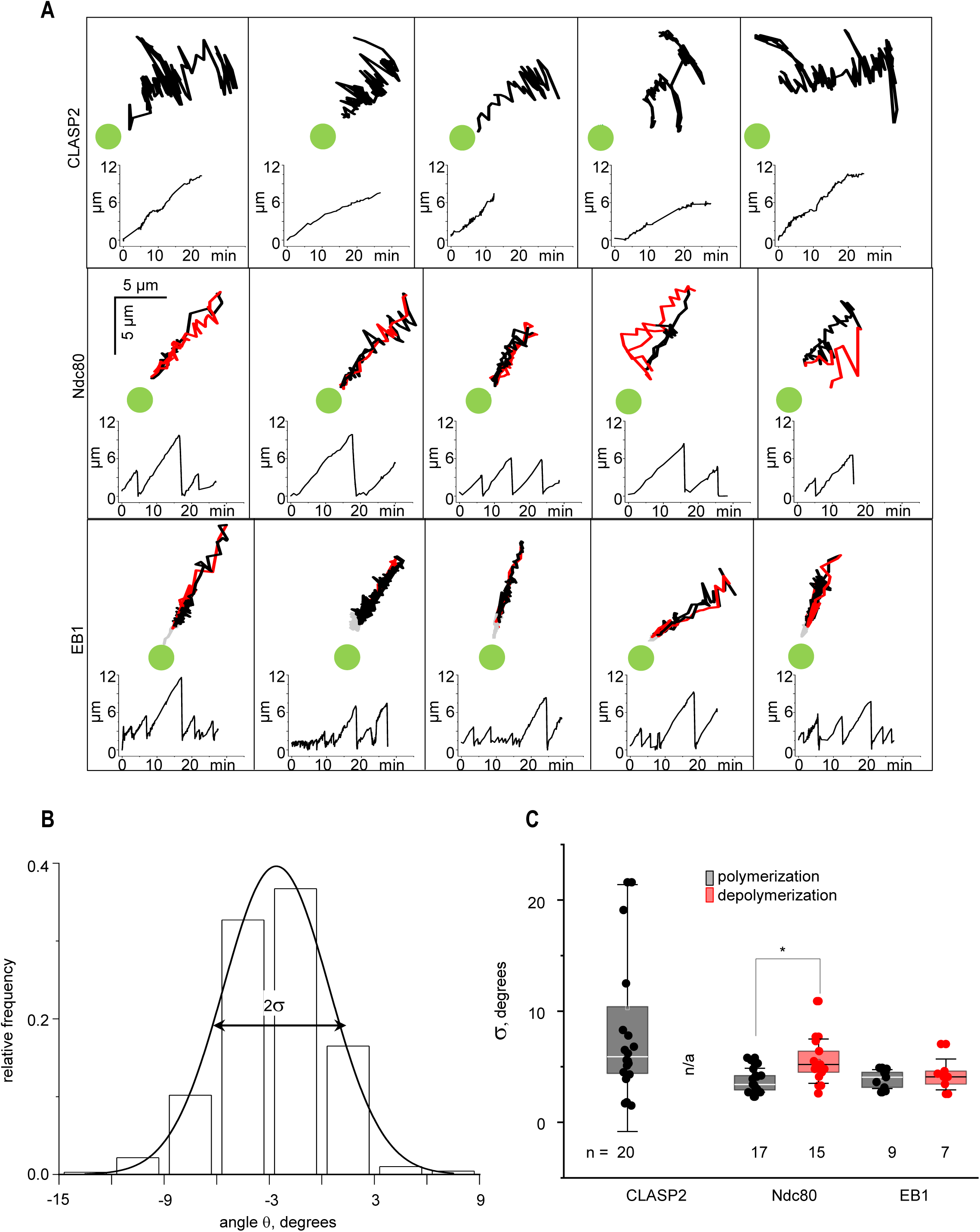
Mechanics of the bead-coupled dynamic microtubule plus-end. **A.** Examples of trajectories and corresponding length vs. time plots for the distal end of Hylite647-labeled microtubule segment, see main text for more details. First panels for Ndc80 and CLASP2α are same as in the main text, but depicting more details about microtubule end dynamics: black parts of the trajectories correspond to the polymerization-coupled movement of microtubule end, and red color for depolymerization. Ndc80 and EB1 proteins support coupling of both elongating and shortening microtubules. **B.** Example histogram of the pivoting angles for one microtubule coupled with CENP-E and EB1. Half-width of the Gaussian fitting, represented with parameter σ, was used as a measure (in degrees) of the directional instability of the coupled ends. **C.** Directional instability (parameter σ) calculated separately for the polymerizing (black) and depolymerizing (red) microtubule ends. During fast disassembly, Ndc80-coupled end slightly softens, presumably because Ndc80 cannot keep up with this end. EB1, which diffuses on microtubule wall much faster, maintains consistently low end rigidity; *p<0.05 (Mann -Whitney U test).

**Supplementary Fig. 10.**
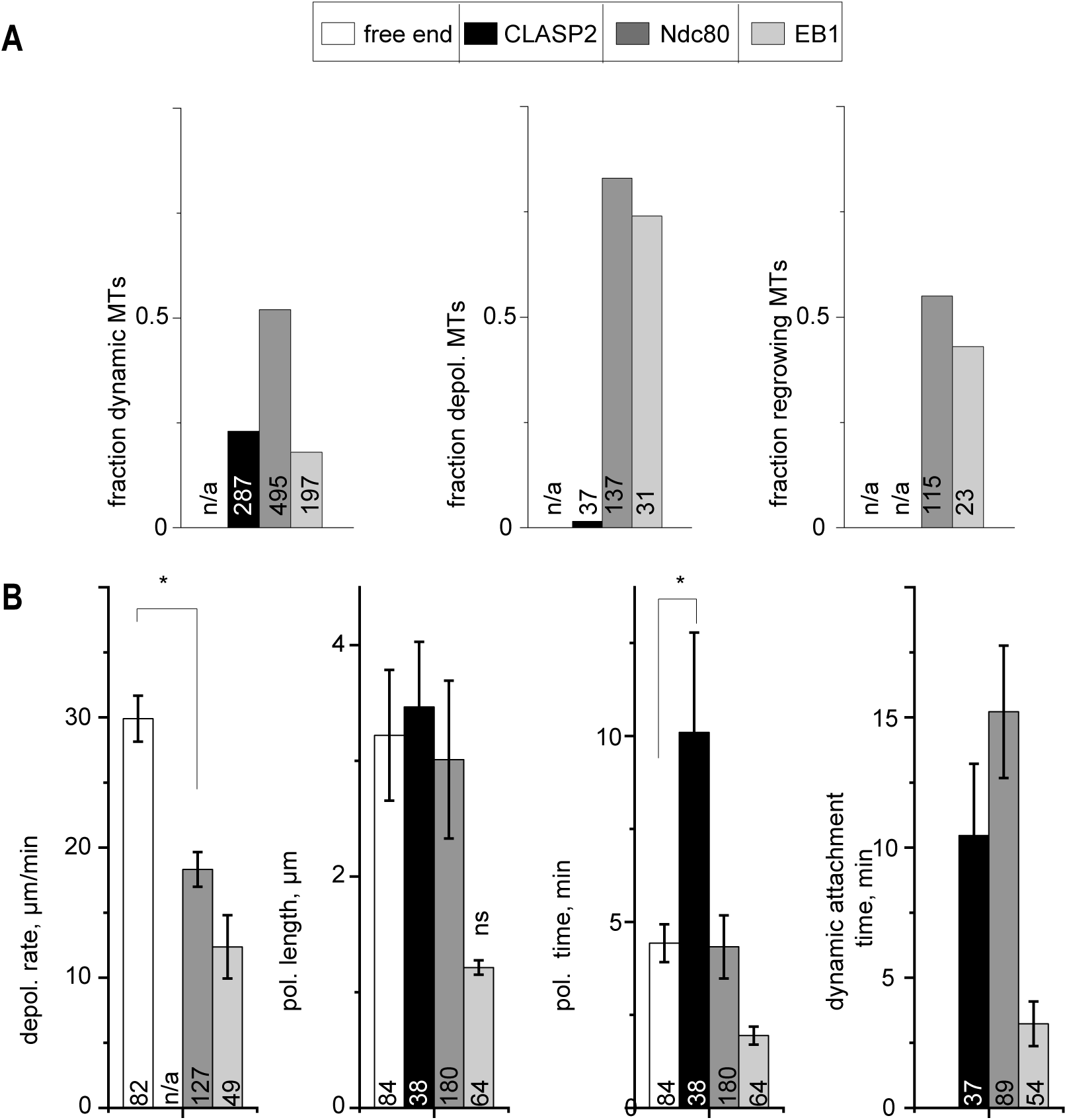
Characterization of the dynamic coupling by beads with different molecular composition. **A.** Frequency of the observed dynamic states for the microtubule ends coupled to beads coated with CENP-E and indicated MAP; data are based on N=4 for Ndc80, N=4 for CLASP2, N=3 for EB1. Fraction of dynamic attachments was calculated from the total number n (given for each column) of bead-bound microtubules. Fraction of microtubules that switched into depolymerization is from the total number n of polymerizing microtubules. Fraction of regrowth events is from the total number n of depolymerizing microtubules. n/a – not applicable. **B.** Dynamic parameters for the freely growing microtubules (N=4) and for the microtubule ends coupled to the protein-coated beads as in panel (b).; n/a – not applicable. **C.** Quantifications for the depolymerization rate (left panel), microtubule polymerization phase (middle two graphs) and total duration of the dynamic microtubule end attachment (last graph); n/a – not applicable. Columns (Mean ± SEM) are calculated based on N≥3 independent experiments and n examined microtubules (noted at each column) *p<0.05 (Mann-Whitney u test).

**Supplementary Fig. 11.**
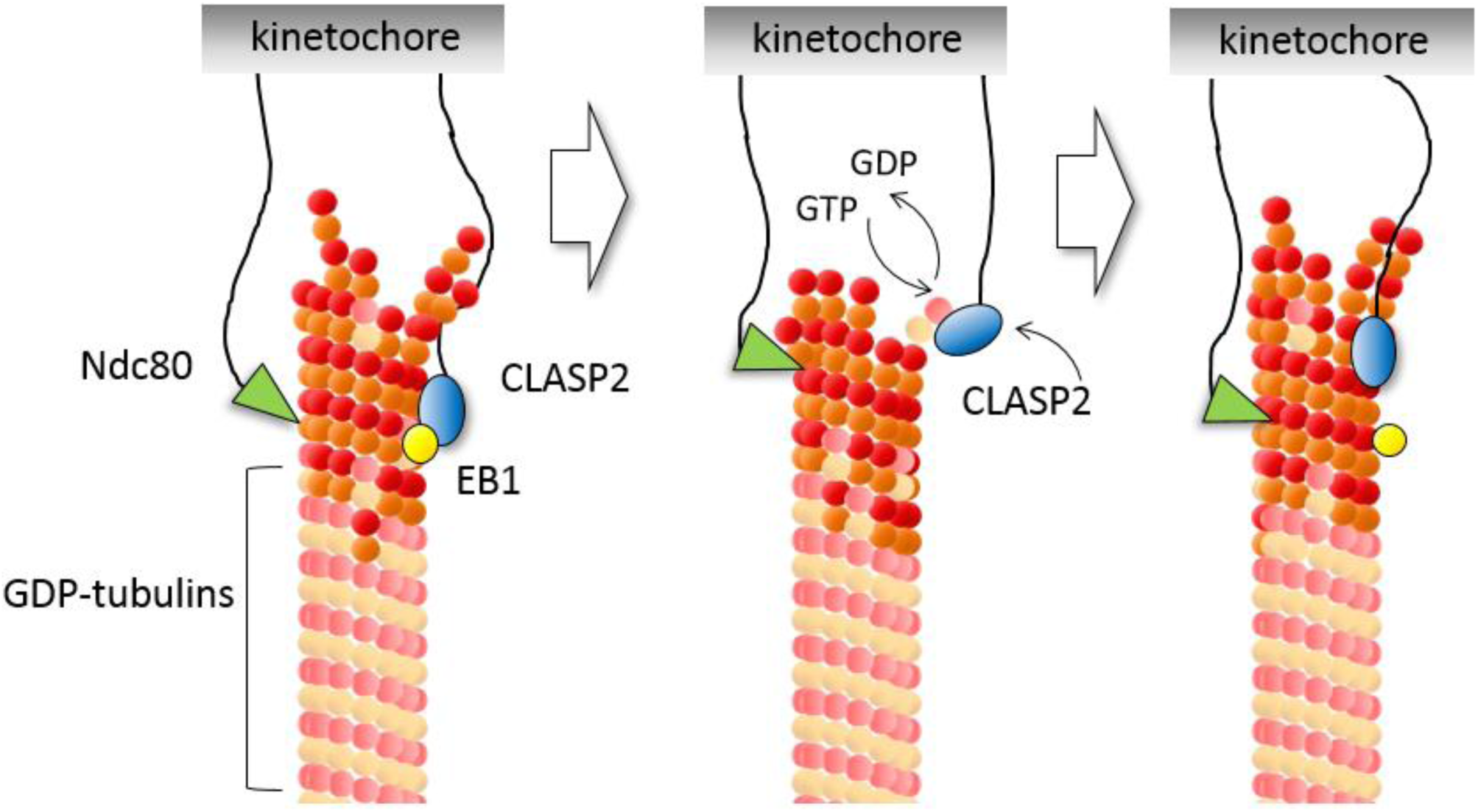
Proposed mechanism of CLASP2-dependent elongation of kinetochore microtubules. Plus-ends of kinetochore microtubules are coupled to kinetochore with the help of numerous proteins, but this simplified schematics depicts only the Ndc80 complex, CLASP2 and soluble EB. CLASP2 is enriched at kinetochore-bound microtubule ends owing to its localization at the kinetochore and via the EB-dependent recruitment. Microtubule wall diffusion of CLASP2 may also increase its chances to encounter the “rogue” tubulins at the microtubule tip. When terminal GDP-tubulin is present, the TOG2 domain of CLASP2 forms a strong bond and stabilizes this dimer temporarily until its associated GDP is replaced with GTP. TOG2 is then released and the protofilament with now terminal GTP-tubulin resumes its normal assembly.

## LEGENDS TO VIDEOS

**Video 1. Microtubule binding to the clusters of CLASP2**

TIRF imaging of the coverslip with immobilized DNA origami (blue), GFP-CLASP2α (green) seen as brighter dots colocalizing with DNA origami and dim dots representing randomly bound molecules, and rhodamine-labeled taxol-stabilized microtubules (red). Microtubules are seen as blurry vibrating spots; most of them remain near the clusters of CLASP2α for the entire imaging time (4.7 hrs). Scale bar 5 µm. See Data source file for more details about this and other videos.

**Video 2. Microtubule binding to the clusters of Ndc80**

Same experiment as in Video 1 but using GFP-tagged Ndc80 Broccoli. Microtubules rotate and slowly diffuse on the Ndc80 clusters during the 2-min imaging sequence. Scale bar 5 µm.

**Video 3. Microtubule binding to the clusters of CLASP2α under flow**

Same experiment as in Video 1 but a buffer is introduced into the chamber flowing from left to right. The removal of unbound microtubules clears the background, whereas the CLASP2α-tethered microtubules orient along the flow. The tethered microtubules start adopting their preferred orthogonal orientation when the flow is stopped later in this sequence, suggesting that the binding sites for CLASP2α are at the very termini of the microtubules. Scale bar 5 µm.

**Video 4. Kinesin-dependent motion of microtubules away from the origami-based clusters of CLASP2α**

Rhodamine-labeled taxol-stabilized microtubules were allowed to bind to GFP-CLASP2α (green) conjugated to origami nano-circles (blue), and the flow was introduced to promote microtubule reorientation and landing on the coverslip coated with Kinesin 1 motor. The video starts when ATP is introduced into the chamber; after a lag, microtubules begin to glide away from the CLASP2α clusters. Scale bar 5 µm.

**Video 5. Unbinding of CLASP2α from the ends of microtubules that incorporate GTP-tubulin.**

This video starts with rhodamine-labeled GMPCPP microtubule seeds (red) bound with their ends to the coverslip-immobilized clusters of GFP-CLASP2α (green). Hilyte647-labeled GTP-tubulin (grey) is then flowed into the chamber to trigger elongation of microtubules from the ends of the GMPCPP seeds. Majority of the microtubule seeds detach immediately and are removed by the flow, although one seed remains bound to origami, presumably due to a stochastic variation in the number and mode of the attached CLASP2α molecules. Subsequently, this seed alternates between two attachment modes to the CLASP2α cluster: the end-on and lateral binding modes. When microtubule seed is in the end-on orientation, the bound end shows no tubulin incorporation, suggesting that CLASP2α binding to the sites at the microtubule tip is not compatible with tubulin incorporation. However, when microtubule orients laterally presumably because some CLASP2α molecules are associated with the wall of this microtubule, its plus-end is freed to add and lose tubulin dimers. Scale bar 5 µm.

**Video 6. Microtubule gliding with CLASP2α in solution.**

This video is paused initially to show that Hilyte647 labeled taxol-stablized microtubules (red) bind GFP-CLASP2α (1 nM, green) preferentially at one of the microtubule ends (white arrows). Additional weaker dots are seen along microtubule walls, at the opposite microtubule ends and also on the coverslip surface. Gliding is driven by the coverslip-associated Kinesin 1, and the brighter microtubule ends are trailing. Scale bar 5 µm.

**Video 7. Dynamic microtubule coupled to CLASP2α-coated bead.**

Video shows a short GMPCPP-stabilized microtubule seed labeled with Hilyte-647 (red) moving away from a coverslip-immobilized 1-μm bead coated with the GFP-labeled CLASP2α and the motor domains of CENP-E kinesin. To ensure visualization of the full range of motions, imaging was done via epi-fluorescence and in the presence of unlabeled soluble tubulin. Continuous motion of the seed away from the bead is associated with new tubulin incorporation at the bead-tethered microtubule plus-end, whereas angular deviations indicate temporary loss of the integrity of the bead-bound microtubule tip. Video plays 30 times faster than actual.

**Video 8. Dynamic microtubule coupled to Ndc80-coated bead.**

This video shows the same experiment as in Video 7 but using GFP-tagged Ndc80 Broccoli instead of CLASP2α (37). Several red microtubule seeds are bound at the bead surface, with one of them moving repeatedly away and toward the bead with no significant pivoting. Video plays 30 times faster than actual. Scale bar 5 µm.

